# Genomic evidence of genetic variation with pleiotropic effects on caterpillar fitness and plant traits in a model legume

**DOI:** 10.1101/518951

**Authors:** Zachariah Gompert, Megan Brady, Farzaneh Chalyavi, Tara C. Saley, Casey S. Philbin, Matthew J. Tucker, Matt L. Forister, Lauren K. Lucas

## Abstract

Plant-insect interactions are ubiquitous, and have been studied intensely because of their relevance to damage and pollination in agricultural plants, and to the ecology and evolution of biodiversity. Variation within species can affect the outcome of these interactions, such as whether an insect successfully develops on a plant species. Whereas specific genes and chemicals that mediate these interactions have been identified, studies of genome-or metabolome-wide intraspecific variation might be necessary to better explain patterns of host-plant use and adaptation often observed in the wild. Here, we present such a study. Specifically, we assess the consequences of genome-wide genetic variation in the model plant *Medicago truncatula* for *Lycaeides melissa* caterpillar growth and survival (i.e., larval performance). Using a rearing experiment and a whole-genome SNP data set (>5 million SNPs), we show that polygenic variation in *M. truncatula* explains 9–41% of the observed variation in caterpillar growth and survival. We detect genetic correlations among caterpillar performance and other plant traits, such as structural defenses and some anonymous chemical features; these genetic correlations demonstrate that multiple *M. truncatula* alleles have pleiotropic effects on plant traits and caterpillar growth or survival (or that there is substantial linkage disequilibrium among loci affecting these traits). We further show that a moderate proportion of the genetic effect of *M. truncatula* alleles on *L. melissa* performance can be explained by the effect of these alleles on the plant traits we measured, especially leaf toughness. Taken together, our results show that intraspecific genetic variation in *M. truncatula* has a substantial effect on the successful development of *L. melissa* caterpillars (i.e., on a plant-insect interaction), and further point toward traits mediating this genetic effect.

## Introduction

Organisms interact with members of other species in myriad ways, including competition for resources, predation, parasitism, herbivory, mutualism and pollination. Phenotypic and genetic variation within species can affect the outcome of these interspecific interactions (Bolnick *et al.*, 2002; Crutsinger *et al.*, 2006; Farkas *et al.*, 2013; Thompson, 2013; Hendry, 2016). For example, a genetic polymorphism for cryptic color pattern affects the probability that *Timema cristinae* stick insects are predated by birds (Nosil, 2004; Nosil *et al.*, 2018), and allelic variation in *Daphnia magna* and its bacterial microparasite, *Pasteuria ramosa,* alters infection rates (Carius *et al.*, 2001; Luijckx *et al.*, 2011, 2013). Intraspecific variation can also affect the establishment and evolution (or co-evolution) of new interactions, including those that form following species introductions (e.g., Cox, 2004; Strauss *et al.*, 2006; Lankau, 2012; Mandeville *et al.*, 2017).

Interactions between plants and herbivorous insects have received considerable scientific attention due to their ubiquity (Forister *et al.*, 2015), their agricultural relevance (Via, 1990; Schoonhoven *et al.*, 2010), and their hypothesized contribution to the extreme biodiversity of these taxonomic groups (via co-evolutionary diversification; Ehrlich & Raven, 1964; Mitter *et al.*, 1988; Fordyce, 2010; Edger *et al.*, 2015; Braga *et al.*, 2018). These interactions are often affected by genetic variation within species, including variation in plant resistance to insects, and for insect acceptance of and performance on potential host plants (e.g., Rausher & Simms, 1989; Via, 1990; Berenbaum & Zangerl, 1998; Stowe, 1998; Dambroski *et al.*, 2005; Ordas *et al.*, 2009; Schoonhoven *et al.*, 2010; Gompert *et al.*, 2015; Mitchell *et al.*, 2016; Nouhaud *et al.*, 2018). Progress in explaining this variation has been made by identifying specific phytochemicals responsible for resistance to insects (e.g., furanocoumarins and glucosinolates), as well as the insect genes and pathways that detoxify these compounds (e.g., cytochrome P450 enzymes, nitrile specifier protein, etc.; Li *et al.*, 2003; Wen *et al.*, 2006; Wheat *et al.*, 2007; Schoonhoven *et al.*, 2010). Genomic and metabolomic approaches have begun to provide a more complete view of how within-species variation affects plant-insect interactions (e.g., Harrison *et al.*, 2018; Nallu *et al.*, 2018). As an example, a recent study of intraspecific variation across 770 traits (including 753 chemical features) in alfalfa showed that among-plant variation in insect herbivore communities was best explained by non-linear interactions among suites of plant traits (Harrison *et al.*, 2018). Such findings highlight the need for quantitative, genome-, phenome-and metabolome-scale analyses of the ecological and evolutionary consequences of intraspecific variation in plant-insect systems. In fact, these approaches may be necessary to explain the geographic mosaic of host-plant use and plant-insect co-evolution found in nature (but see, e.g., Berenbaum & Zangerl, 1998), in other words, to address questions such as: (i) Why are certain plant species fed on by a species of insect in some places but not others?, and (ii) To what extent do different host-plant populations represent distinct adaptive landscapes?.

Here, we take an initial step towards this larger aim by quantifying the effect of genome-wide plant genetic variation on caterpillar performance (weight and survival) in the Melissa blue butterfly, *Lycaeides melissa* (Lepidoptera: Lycaenidae). *Lycaeides melissa* butterflies are found throughout western North America where they feed on various legume hosts, particularly from the genera *Astragalus* and *Lupinus* (Scott, 1986). *Medicago sativa* (alfalfa) is a legume native to Eurasia that was introduced to North America ~250 years ago as a forage crop (Michaud *et al.*, 1988). Since then, *L. melissa* has repeatedly colonized *M. sativa,* and numerous *L. melissa* populations now use this plant as their primary host, especially where *M. sativa* has escaped from cultivation along roadsides and trails (Chaturvedi *et al.*, 2018). *Lycaeides melissa* populations that use *M. sativa* show evidence of adaptation to this host, such as increased oviposition preference and larval performance (Forister *et al.*, 2012; Gompert *et al.*, 2015). However, *M. sativa* remains an inferior host in terms of laraval performance relative to other common hosts, and many *M. sativa* populations are not used by *L. melissa* within *L. melissa*’s range (Forister *et al.*, 2009). Thus, host use in *L. melissa* comprises a mosaic of occupied and unoccupied patches of *M. sativa* and native legume hosts. Previous experiments documented genetic variation within *L. melissa* populations for larval performance on *M. sativa* (Gompert *et al.*, 2015), and also showed that *M. sativa* populations vary in their suitability for *L. melissa* caterpillars (Harrison *et al.*, 2016). However, past experiments were not designed to parse genetic versus environmental contributions to host-plant suitability (this distinction is critical for co-evolutionary dynamics), nor to identify specific plant traits (or plant genes) affecting *L. melissa* caterpillar performance.

Our ultimate goal is to explain variation in the (relatively recently established) interaction between *M. sativa* and *L. melissa* across the landscape. This includes determining to what extent genetic differences among *M. sativa* plants affect whether a *M. sativa* population is colonized by *L. melissa,* and to what extent genetic differences among plant populations affect subsequent ecological and evolutionary dynamics and outcomes (e.g., *L. melissa* demographics, the degree of host adaptation, etc.). Despite its role in agriculture, genomic resources for *M. sativa* are limited. Consequently, in the present study we use the model plant *Medicago truncatula* as a proxy for *M. sativa. Medicago truncatula* is a close relative of *M. sativa* that occurs throughout the Mediterranean basin in Europe and is cultivated in Australia (Choi *et al.*, 2004a, b). Because of its modest genome size (~500 million base pairs), simple diploid genetics, and short generation time (~10 weeks), *M. truncatula* has been developed as the model species for legumes (Young & Udvardi, 2009; Young *et al.*, 2011). Resources for this species include a high-quality reference genome and hundreds of fully sequenced, inbred lines derived from natural accessions (Young *et al.*, 2011; Stanton-Geddes *et al.*, 2013). Unlike *M. sativa, M. truncatula* is not found in North America and thus is not available as a host for *L. melissa* (i.e., it is not part of *L. melissa*’s realized niche). However, both *Medicago* species could be used by other *Lycaeides* in Eurasia where most of the biodiversity in this genus is found (North American *Lycaeides* are descended from Eurasian ancestors that came across the Bering land bridge about two million years ago; Gompert *et al.*, 2008; Vila *et al.*, 2011). Thus, while our results do not directly assess variation in *M. sativa*, they can show the potential for intraspecific plant genetic variation to affect plant-insect interactions in this system; further, we hypothesize that the *M. trunatula* genes and traits affecting caterpillar performance will function similarly in *M. sativa*.

In this study, we combine statistical genomic methods with a caterpillar rearing experiment to assess the effect of *M. truncatula* phenotypic and genetic variation on *L. melissa* caterpillar performance. We address the following specific questions: (i) How much of the variation in *L. melissa* growth and survival can be explained by genetic variation in *M. truncatula*?, (ii) Do genetic loci that affect a set of measured plant traits (some putatively associated with plant vigor or defense) have pleiotropic effects on caterpillar performance?, and (iii) How well do the effects of *M. truncatula* alleles on the measured plant traits explain their effects on caterpillar performance. Thus, we quantify the direct effect of *M. truncatula* genetic variation on caterpillar performance, and its effect through a set of plant traits. We think that this combination of approaches has the potential to (a) provide a more mechanistic understanding of this plant-insect interaction by connecting genetic patterns with plant traits, and (b) discover previously unhypothesized sources of variation in caterpillar performance by identifying alleles associated with caterpillar performance that are not associated with any of the plant traits we measured. Moreover, the methods and approaches we use allow us to generate statistical and functional information about the genetic basis of this interaction even if it is polygenic (see Methods and Results for details).

## Methods

### Plant propagation and trait measurements

We obtained seeds from 100 *M. truncatula* lines, which are part of the *Medicago* HapMap project (http://www.medicagohapmap.org). Seeds (i.e., germplasm) were obtained from INRA-Montpellier (Montpellier, France), and from the USDA Agricultural Research Station at Washington State University (Pullman, WA, USA; Table S1). Each line was derived from a natural accession, but has since been inbred to near complete homozygosity. Whole genome sequences are available for each line (Branca *et al.*, 2011; Stanton-Geddes *et al.*, 2013), and the lines have been used in other genome-wide association mapping studies (GWAS), including GWAS on biomass, drought-related traits, plant defenses, flowering time, and nodulation (e.g., Stanton-Geddes *et al.*, 2013; Kang *et al.*, 2015).

**Table 1:** Plant traits along with our predictions about their primary functional roles and relationships with caterpillar performance. We are presenting simplified predictions to guide interpretation, but are aware that the traits potentially have multifaceted relationships to growth and defense. ↗ denotes a positive correlation with caterpillar performance, whereas ↘ denotes a negative relationship with caterpillar performance. Our classification of SLA is based on its general association with mechanical properties of leaves, including work to shear, tear and penetrate (reviewed in Hanley *et al.*, 2007). All 19 IR chemical features are treated together here, and thus we predict that they include a mixture of features associated with vigor (↗) and defense (↘). Leaf shape is not included in the table, as its putative function and effects are not known.

| Traits | Primary putative function | Predicted relationship with caterpillar performance |
| --- | --- | --- |
| Leaf length | Growth | ↗ |
| Leaf width | Growth | ↗ |
| Leaf area | Growth | ↗ |
| Leaf weight | Growth | ↗ |
| SLA | Defense | ↘ |
| Trichome den. | Defense | ↘ |
| Leaf tough. | Defense | ↘ |
| Plant height | Growth | ↗ |
| IR features | Growth or defense | ↗ or ↘ |

We planted five replicate pots with seeds from each of the 100 *M. truncatula* lines on May 4th and 5th, 2017 (see “Planting and tending *Medicago truncatula*” in the online supplemental material [OSM] for additional details). *Medicago truncatula* plants were grown in a greenhouse under ambient light (~14–15 hours of daylight) at approximately 18–27°C (with variable humidity), and were watered daily or every other day as needed. We thinned the *M. truncatula* seedlings on May 26th (i.e., after germination was complete) to ensure that no pots had more than two plants. This was done to minimize competition among plants, while still providing sufficient plant biomass for the caterpillar rearing experiments. A few plant lines had low germination rates and were dropped from the experiment leaving us with 94 lines, each with five replicate pots.

We measured a series of morphological traits potentially associated with plant vigor or resistance to insects (e.g., putative structural plant defenses; Table 1; Levin, 1973; Hanley *et al.*, 2007; Malishev & Sanson, 2015). First, 20 days after planting, we measured leaf size (length, width and area), **leaf shape** (length/width), **trichome density, dry leaf weight** and **specific leaf area** (SLA) for each plant line and replicate (pot) (we haphazardly selected one of the two plants in each pot for taking measurements). We chose the second true leaf for these measurements (that is leaf 1 from branch B0, see Figs. 1 & 2 from Moreau, 2006). We measured the width (at the widest point) and length (along the midvein) of the middle leaflet with calipers (each leaf comprises three leaflets; measurements were taken to the nearest 1 mm). Next, we calculated leaf area (length × width) and shape (length/width) from these measurements. We then counted the number of trichomes in a 2.5 mm diameter circle directly adjacent to the midvein under a stereoscope (35× magnification). The three leaflets from each plant were then placed in a coin envelope in a bin with desiccant. The dry weight of the middle leaflet from each of these leaves was measured on a Mettler Toledo XPE105 analytical microbalance (Mettler Toledo) to the nearest 0.01 mg. Leaf area and dry weight were used to calculate SLA (SLA is the ratio of leaf area to dry mass and is often correlated with leaf mechanical properties, such as work to tear, shear or punch; Hanley *et al.*, 2007).

**Figure 1:**
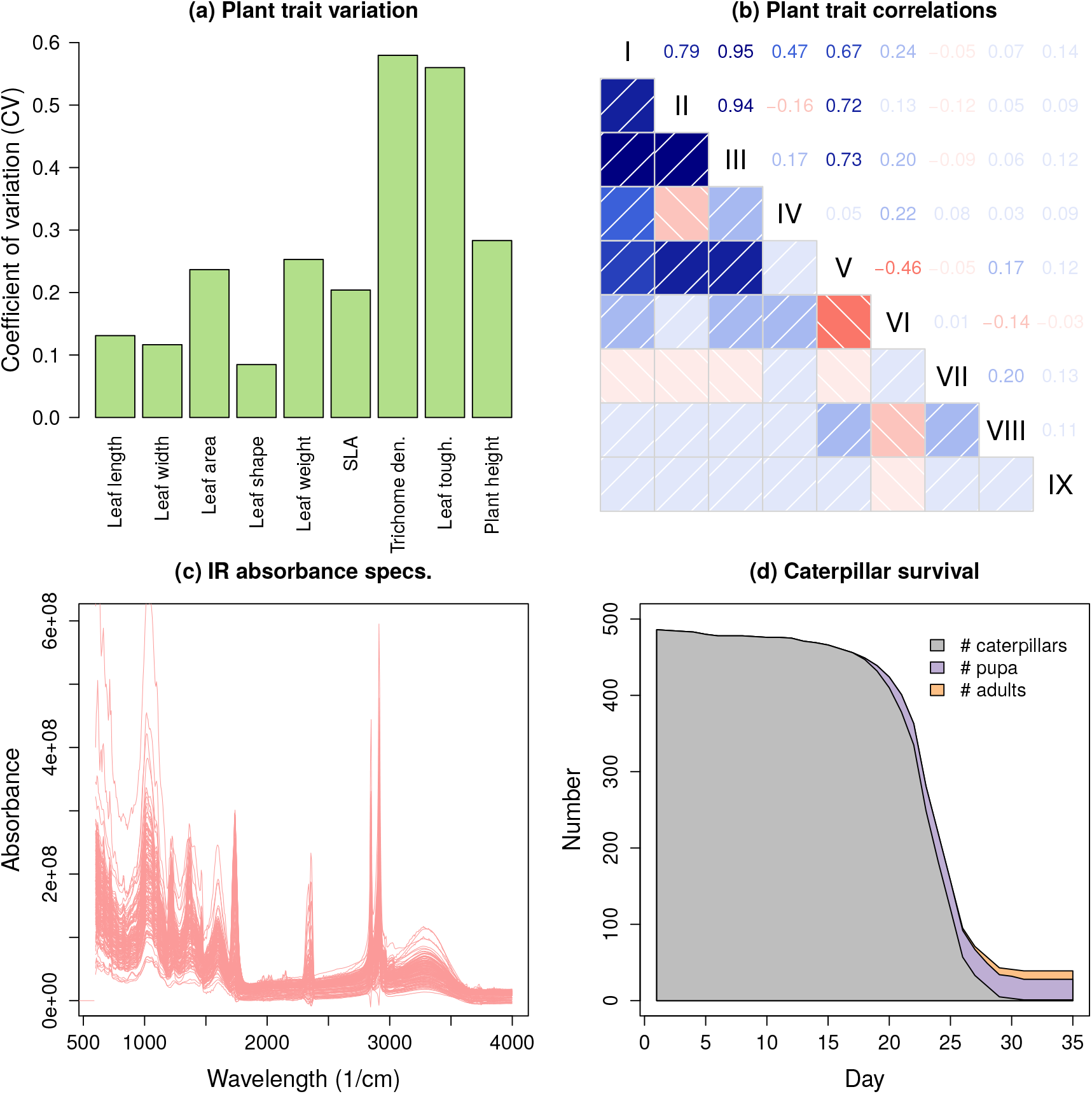
Plant (*M. truncatula*) and caterpillar (*L. melissa*) phenotypic data. Panel (a) provides the coefficient of variation (CV) for each of the nine plant growth/defense traits. Panel (b) presents pairwise phenotypic correlations for the same nine traits (roman numerals denote the trait numbers ordered as in panel a). Pearson correlations are shown in the upper triangle of the correlation matrix, and depicted graphically in the lower triangle of the correlation matrix, with darker shading denoting higher correlations. Panel (c) shows the infrared (IR) absorbance spectra for each plant (one line per plant) (see Fig. S2 for phenotypic correlations for the IR traits). Panel (d) gives the number of caterpillars, pupa, and adults (and thus the total number of *L. melissa*) alive at 0 to 35 days of age.

**Figure 2:**
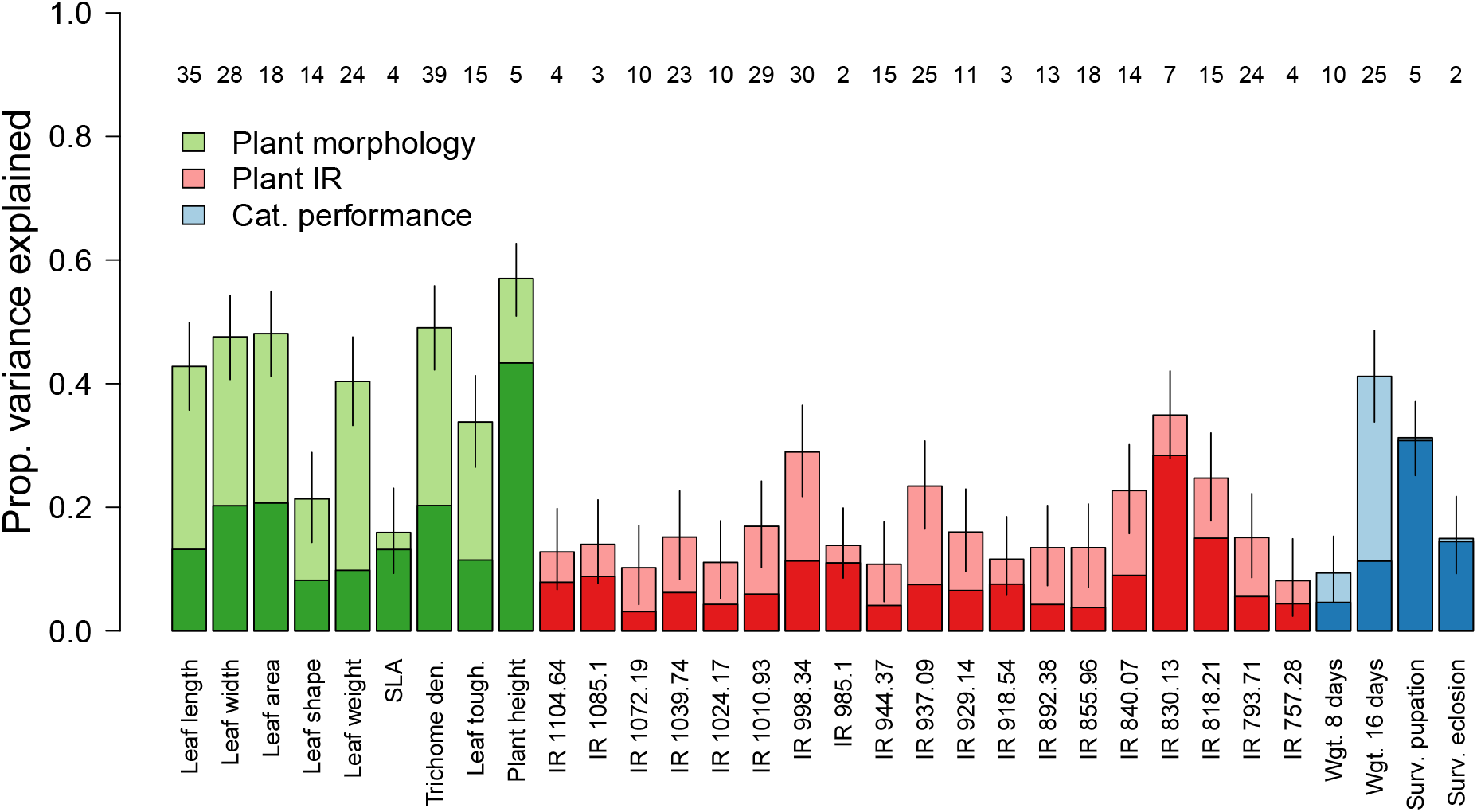
Graphical summary of plant (*M. truncatula*) and caterpillar (*L. melissa*) trait variation explained by *M. truncatula* genetics. Bars denote the posterior median for the proportion of trait variation explained by plant genetics (PVE); vertical lines denote the 90% equal-tail probability intervals (ETPIs). Darker shaded regions of the bars provide point estimates (posterior median) for the subset of the PVE attributed to genetic variants with measurable effects (as opposed to infinitesimal effects). Numbers along the top of the plot give point estimates (posterior median) for the number of causal variants affecting each trait (i.e., total number of distinct QTL). See Table S2 for detailed quantitative summaries of these parameter estimates, including measures of uncertainty in each parameter.

**Table 2:** REML estimates for each trait of the proportion of phenotypic variation found among *M. truncatula* lines (‘Prop. var.’). Test statistics (LR = likelihood ratios) and *P*-values from the null hypothesis test of no line effect are reported.

| Traits | Prop. var. | LR | *P* |
| --- | --- | --- | --- |
| Leaf length | 0.43 | 115.87 | <0.001 |
| Leaf width | 0.49 | 144.78 | <0.001 |
| Leaf area | 0.49 | 147.90 | <0.001 |
| Leaf shape | 0.21 | 32.23 | <0.001 |
| Leaf weight | 0.41 | 102.49 | <0.001 |
| SLA | 0.15 | 17.12 | <0.001 |
| Trichome den. | 0.49 | 151.82 | <0.001 |
| Leaf tough. | 0.34 | 69.95 | <0.001 |
| Plant height | 0.59 | 218.92 | <0.001 |
| IR 1104.64 | 0.13 | 12.79 | <0.001 |
| IR 1085.1 | 0.15 | 16.20 | <0.001 |
| IR 1072.19 | 0.10 | 7.01 | 0.004 |
| IR 1039.74 | 0.16 | 17.46 | <0.001 |
| IR 1024.17 | 0.10 | 8.48 | 0.001 |
| IR 1010.93 | 0.17 | 20.27 | <0.001 |
| IR 998.34 | 0.29 | 54.85 | <0.001 |
| IR 985.1 | 0.12 | 10.78 | <0.001 |
| IR 944.37 | 0.10 | 7.46 | 0.003 |
| IR 937.09 | 0.23 | 35.98 | <0.001 |
| IR 929.14 | 0.15 | 16.76 | <0.001 |
| IR 918.54 | 0.12 | 11.63 | <0.001 |
| IR 892.38 | 0.13 | 11.99 | <0.001 |
| IR 855.96 | 0.13 | 13.19 | <0.001 |
| IR 840.07 | 0.23 | 36.36 | <0.001 |
| IR 830.13 | 0.36 | 80.73 | <0.001 |
| IR 818.21 | 0.24 | 40.84 | <0.001 |
| IR 793.71 | 0.14 | 15.10 | <0.001 |
| IR 757.28 | 0.09 | 6.05 | 0.007 |
| Wgt. 8 days | 0.07 | 3.99 | 0.019 |
| Wgt. 16 days | 0.41 | 93.69 | <0.001 |
| Surv. pupation | 0.24 | 40.33 | <0.001 |
| Surv. eclosion | 0.05 | 2.39 | 0.059 |

We measured **plant height**, from the cotyledons to the tip of the **longest branch**, 31 days after planting (again, we haphazardly selected one of the two plants in each pot for taking this measurement). Leaf toughness was measured 33 days after planting using a penetrometer. We selected the main leaf from the second primary branch for this assay. The force required to penetrate each of the three leaflets along the midvein was recorded. We took the mean of these three measures as a metric of leaf toughness.

Plant chemistry was quantified with **attenuated total reflectance infrared** (ATR-IR) spectroscopy. ATR-IR spectroscopy constitutes a quick, cost-effective method to analyze a range of organic chemical compounds in plant and animal tissues. Although the absorbance is directly related to the concentration of specific chemical signatures, there is not a simple one-to-one relationship between IR spectral patterns and specific chemical compounds of interest. Moreover, spectral features are the summation of similar overlapping IR transitions, representative of various compounds within a tissue. Consequently, IR data are often combined with more specific compositional analyses (e.g., HPLC-MS). The combined data can be used to construct a multivariate model linking IR spectral data to chemical compounds (e.g., Foley *et al.*, 1998; Ramirez *et al.*, 2015; Costa *et al.*, 2018). This was not our goal here. We instead used IR spectral features as anonymous chemical markers (akin to AFLPs for genetic analyses) which could be connected to the presence of specific molecules in future work using compositional methods such as liquid chromatography–mass spectrometry.

Infrared spectra were collected using a Thermo Nicolet 6700 FTIR (a high-resolution instrument with a diamond crystal ATR), which was used to scan 4000-600 cm^−1^ of the infrared spectrum. Leaves were placed in direct contact with the diamond crystal, and the average of 32 scans was recorded for each leaf surface with 4 cm^−1^ resolution. A Norris-Williams second derivative spectrum was calculated for each transmittance measurement using 5-point smoothing and a gap size of 5 segments (absorbance is directly proportional to concentration [Beer’s Law], and absorbance =-log(transmittance)). We focused on the subset of IR features between ~750 and 1100 cm^−1^ and with >10% of the phenotypic variation partitioned among plant lines (see Fig. S1).

### Caterpillar husbandry and performance assays

We obtained neonate *L. melissa* caterpillars for larval performance assays on the *M. truncatula* accessions. First, 26 female *L. melissa* butterflies were collected on June 5th (2017) from a site along the Bonneville shoreline trail in northern Utah, USA (41.725°N, 111.794°W, 1513 m elevation). As in past work (e.g., Forister *et al.*, 2013; Gompert *et al.*, 2015), these butterflies were caged individually in plastic oviposition chambers along with a few sprigs of their host plant (*Medicago sativa*). After 48 hours, *L. melissa* eggs were collected from the host-plant material and placed in unvented Petri dishes in a Percival incubator (model no. 136VL; 27°C; 14 hrs. light:10 hrs. dark) until they hatched.

Caterpillars began to emerge on June 9th, and were then placed in individual unvented Petri dishes with a leaf from one of the 94 *M. truncatula* accessions (i.e., on one of the 94 plant lines). We inspected caterpillars daily, adding new leaf material from the same plant line as needed (as in Gompert *et al.*, 2015). We rotated the replicate/pot used for each plant line each day. Thus, caterpillars only ate leaves from a single plant line (genotype), but fed on all five replicate pots. Caterpillars were maintained in a Percival incubator at 27°C with 14 hour days (10 hours of dark). We reared 486 caterpillars total (~5 per plant line). We checked all caterpillars daily for survival and recorded **survival to pupation** and **survival to eclosion** as adults. As an additional metric of performance, we measured **8-and 16-day caterpillar weight** (*L. melissa* caterpillars generally spend 20 to 30 days as larvae) on a Mettler Toledo XPE105 analytical microbalance (Mettler Toledo; weights were recorded to the nearest 0.01 mg). Weight and lifetime fecundity are highly correlated in *L. melissa* (Forister *et al.*, 2009).

### Variance partitioning

Our analyses focus on the 9 plant morphological traits (leaf length, leaf width, leaf area, leaf shape, leaf dry weight, SLA, trichome density, leaf toughness and plant height), 19 IR traits (i.e., anonymous chemical features), and four caterpillar performance traits (weight at 8 days, weight at 16 days, survival to pupation, and survival to eclosion; survival is a binary trait; Table 1). Prior to genetic mapping and genomic prediction, we first quantified the proportion of trait variation found among plant lines (i.e., genotypes) for each of these 32 traits. As we are working with replicated, inbred lines, these are estimates of the broad-sense heritability for each of the traits (with respect to plant not caterpillar genotypes; because caterpillars fed across plants of a genotype, these estimates are upper bounds for the broad-sense heritabilities of the caterpillar performance traits).

We estimated the among-line variance for each trait by fitting linear mixed-effect models via restricted maximum likelihood (REML). This was done with the lmer function in `lme4 R` package (package version 1.1.19, `R` version 3.4.4; Bates *et al.*, 2015). We then tested the null hypothesis that the among-line variance was 0 using an exact restricted likelihood ratio test, which was based on 10,000 simulated values to approximate the null distribution (Crainiceanu & Ruppert, 2004; Greven *et al.*, 2008). This was done with the `exactRLRT` function in the `RLRsim` package in `R` (version 3.1.3; Scheipl *et al.*, 2008).

### *Medicago truncatula* genomic data

Whole-genome SNP data for the *M. truncatula* accessions were obtained from the *M. truncat-ula* HapMap project (http://www.medicagohapmap.org/; version Mt4.01; Stanton-Geddes *et al.*, 2013). These data comprised 40 million SNPs, which were mapped to the *M. truncatula* reference genome v4.0 (we used the quality-filtered SNP `bcf` files; Young *et al.*, 2011). We applied additional quality filters to these data with `vcftools` (version 0.1.15; Danecek *et al.*, 2011) such that we only retained bi-allelic SNPs with minor allele frequencies ≥ 0.01, and with a minimum sequencing depth of 2× per individual, no more than 20% missing data (across the 94 lines analyzed in this study), and a phred-scaled quality score of ≥ 30. We only considered SNPs mapped to the eight *M. truncatula* chromosomes. Approximately 13 million SNPs passed these filters. We then used `plink` (version 1.09; Purcell *et al.*, 2007) to remove redundant SNPs, that is SNPs that were in very high linkage disequilibrium (LD) with each other. Specifically, using the `indep-pairwise` command, one of each pair of high-LD SNPs, defined as r^2^ ≥ 0.8 in a 10 kilobase (kb) window, was pruned. After this step, we retained 5,648,722 SNPs for downstream analyses.

The *M. truncatula* HapMap data set included SNP genotype calls and relative genotype likelihoods generated by `GATK` (McKenna *et al.*, 2010). Rather than use the raw genotype calls (which ignore uncertainty in genotypes and information from population allele frequencies), we used an empirical Bayesian approach to obtain estimates of genotypes based on the genotype likelihoods and a prior defined by the allele frequencies at each locus. As in past work (e.g., Gompert *et al.*, 2015), we first used an expectation-maximization algorithm to obtain maximum likelihood estimates of the allele frequencies for each SNP. This was done with the computer program `estpEM` (in Dryad repository, doi:https://doi.org/10.5061/dryad.nq67q; Soria-Carrasco *et al.*, 2014; Riesch *et al.*, 2017). This program implements the EM algorithm from Li *et al.* (2009) and provides allele frequency estimates that account for genotype uncertainty. Prior probabilities for each genotype were then specified based on the allele frequencies, such that *Pr*(*g_ij_*|*p*_*i*_) ~ binomial(*p_i_*, *n* = 2), where *g_ij_* denotes the genotype at locus *i* for individual *j*, and *p_i_* denotes the non-reference allele frequency. Next, we computed the posterior probability of each genotype according to Bayes theorem, and obtained point estimates (posterior means) for genotypes 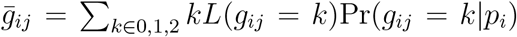, where *L*(*g_j_* = *k*) is the relative genotype likelihood based on the sequence data and associated quality scores. These genotype estimates take on values between 0 (reference-allele homozygote) and 2 (non-reference-allele homozygote), but are not constrained to be integer values.

### Genome-wide association mapping and genomic prediction

We fit Bayesian sparse linear mixed models (BSLMMs; Zhou *et al.*, 2013) with `gemma` (version 0.94.1) to quantify the contribution of *M. truncatula* (i.e., plant) genetic variation to phenotypic variation in the plant traits and *L. melissa* caterpillar performance. Unlike traditional genome-wide association mapping methods, BSLMMs fit a single model with all SNPs simultaneously and thus mostly avoid issues related to testing large numbers of null hypotheses. In particular, trait values are modeled as a function of a polygenic term and a vector of the (possible) measurable effects (associations) of each SNP on the trait (*β*; Zhou *et al.*, 2013). Variable selection is used to estimate the SNP effects; SNPs can be assigned an effect of 0 (not in the model) or a non-zero effect (in the model) (Guan & Stephens, 2011). A Markov chain Monte Carlo (MCMC) algorithm is used to infer the posterior inclusion probability (PIP) for each SNP, that is, the probability that each SNP has a non-zero effect. The polygenic term defines an individual’s expected deviation from the grand phenotypic mean based on all of the SNPs. It accounts for phenotypic covariances among individuals caused by their relatedness or overall genetic similarity (i.e., observed kinship; Zhou *et al.*, 2013). The kinship matrix also serves to control for population structure and relatedness when estimating the effects of individual SNPs (β) along with their PIPs. Likewise, SNPs in LD with the same causal variant effectively account for each other, such that only one or the other is needed in the model, and this is captured by the PIPs.

The hierarchical structure of the model provides a way to estimate additional parameters that describe aspects of a trait’s genetic architecture (Guan & Stephens, 2011; Zhou *et al.*, 2013; Lucas *et al.*, 2018). These include the proportion of the phenotypic variance explained (PVE) by additive genetic effects (this includes β and the polygenic term, and should approach the narrow-sense heritability), the proportion of the PVE due to SNPs with measurable effects or associations (this is called PGE and is based only on β), and the number of SNPs with measurable associations (n-γ). All of these metrics integrate (via MCMC) over uncertainty in the effects of individual SNPs, including whether these are non-zero. Likewise, BSLMMs can be used to obtain genomic estimated breeding values (GEBVs), that is, the expected trait value for an individual from the additive effects of their genes as captured by both β and the polygenic term (Lucas *et al.*, 2018). Most other genomic prediction methods provide GEBVs based solely on a polygenic term (e.g., Meuwissen *et al.*, 2001; Hayes *et al.*, 2009; Ober *et al.*, 2012).

We fit BSLMMs for each of 32 traits using `gemma` (version 0.94.1; Zhou *et al.*, 2013) with 15 MCMC chains each with a 500,000 iteration burn-in followed by 2 million sampling iterations with a thinning interval of 20. GEBVs were obtained using the -`predict 1` option, with predictions averaged over the 15 MCMC chains. GEBVs were used to estimate genetic correlations among traits (i.e., a standardized G-matrix). As a guard against statistical artifacts, we fit BSLMMs to 12 pseudo (randomized)-data sets derived from the caterpillar data (while these methods have been assessed in detail elsewhere, e.g., Zhou *et al.*, 2013; Gompert *et al.*, 2017, we were particularly concerned that the low number of survivors and binary data for survival could lead to spurious association; for details, see “BSLMMs fit to randomized data” in the OSM).

### Connecting plant trait genetics with caterpillar performance

Genetic covariances (correlations) among plant and caterpillar traits (as captured by the G-matrix) can provide evidence of a shared genetic basis for these traits. However, these treat pairs of traits independently and do not formally quantify the total contribution of alleles affecting the measured plant traits to the alleles affecting caterpillar performance. Thus, we next assessed the extent to which we could explain variation in the caterpillar performance GEBVs based on the GEBVs for the plant morphology and chemistry traits, as well as which plant trait GEBVs were most important for this. In other words, we wanted to know how well we could explain (or predict) the caterpillar performance GEBVs (that is, the expected performance trait values based on plant genetics) from the subset of genetic variants associated with phenotyped plant traits (as captured by the plant trait GEBVs, and thus weighted by their effects on the plant traits). High explanatory (or predictive) power would imply that most of the *M. truncatula* genetic variants affecting caterpillar performance either had pleiotropic effects on some of the plant traits we measured or were tightly linked to genetic variants that affected these traits. This should also allow us to identify specific plant traits that share a common genetic basis with (and thus potential causal link to) caterpillar performance. We used two complementary approaches to answer this question: (i) multiple regression with Bayesian model averaging, and (ii) random forest regression. A key distinction between these methods is whether they assume linear (multiple regression) or non-linear (random forest regression) relationships between predictors and response variables. Note that for each plant and caterpillar trait, there was a single GEBV estimate per plant line, and thus the sample size for these analyses was *N* = 94 plant lines.

We used multiple regression with Bayesian model averaging to identify the subset of predictors (plant GEBVs) that best explained variation in caterpillar performance GEBVs, while accounting for uncertainty in the effects of each covariate including which covariates have non-zero effects. The multiple regression models were fit with the `BMS R` package (package version 0.3.4, `R` version 3.4.2; Zeugner & Feldkircher, 2015). Zellner’s g-prior was used for the regression coefficients with *g* = *N*, where *N* is the number of observations (*N* = 94; Zellner, 1986), and a uniform prior was used for the different models (i.e., sets of covariates with non-zero effects; Zeugner & Feldkircher, 2015). Parameter estimates were obtained using MCMC with a 5000 iteration burnin-in and 100,000 sampling iterations, and using the birth-death sampler for exploring model space. We then used 10-fold cross-validation to assess the predictive power of these models (that is, the power of the model to explain observations not used in fitting the model). Predictive power necessarily averages over uncertainty in covariate effects (including which covariates have non-zero effects), and was measured as the Pearson correlation (and squared Pearson correlation) between the observed and predicted caterpillar performance GEBVs. As a simpler metric of explanatory power (not predictive power), we estimated the coefficient of determination (*r*^2^) from a standard linear model that included only the subset of predictors (i.e., plant trait GEBVs) with posterior inclusions probabilities (PIPs) greater than 0.5 in the Bayesian model averaging analysis (importantly, here the same data were used to fit the model and assess its explanatory power). This was done with the `lm` function in `R`.

The random forest regression algorithm was similarly used to determine the influence of the plant trait GEBVs on the caterpillar performance GEBVs, while allowing for non-linear interactions among variables (Breiman, 2001). Random forest creates multiple regression trees and then outputs the importance of each predictor. The number of trees created was left at the default of 500, after determining that changing the number of trees from this number did not significantly reduce error. The number of variables randomly sampled at each split (`mtry`) and the number of terminal nodes (`nodesize`) were chosen to minimize OOB error by manually varying these parameters from one to 20 (all possible combinations were considered). To determine variable importance, the predictor of interest was varied and the percent change mean-squared error (%MSE) in predicting the out-of-bag (OOB) data was determined for each. Those with the greatest effect on %MSE are the most important predictor variables. Random Forest was run using `randomForest` package (version 4.6-12) in `R` (Liaw & Wiener, 2002). Random forest regression was `run` separately with each of the caterpillar performance GEBVs as the response and the GEBVs for plant traits as predictors.

## Results

### Variation in plant traits and caterpillar performance

We documented substantial phenotypic variation for all 32 traits assayed (e.g., Fig. 1a,c). Phenotypic correlations among traits were evident, particularly among leaf morphology traits (some of which are functions of each other; Fig. 1b) and among some IR chemical traits (Fig. S2). Caterpillar survival rates were initially high, with only nine of the 486 caterpillars (1.9%) dying within the first eight days; the mean survival time was 22.3 days (excluding caterpillars that pupated; Fig. 1d). But most caterpillars failed to pupate (448, or 92.2%), such that high mortality rates were observed between 20 and 30 days of larval development. Of the 38 caterpillars that did pupate, 11 eclosed as adults (29%) (several of the adults were deformed). Mean caterpillar weight at 8 and 16 days were 5.1 mg (s.d. = 2.5 mg, min. = 0.04 mg, max. = 12.9 mg) and 17.7 mg (s.d. = 7.7 mg, min. = 3.02 mg, max. = 82.7 mg), respectively.

The 32 traits exhibited significant among-line variation, with the possible exception of survival to eclosion as adults (Table 2). The proportion of variation among lines ranged from 0.15 (SLA) to 0.59 (plant height) for the plant morphology traits, from 0.09 to 0.36 for the plant IR traits, and from 0.05 (survival to eclosion) to 0.41 (16 day weight) for the caterpillar performance traits (Fig. S3). With the exception of survival to eclosion (restricted likelihood ratio test [RLRT], P = 0.059), the null model of no among-line variance could be confidently rejected for all traits (RLRT, all *P* < 0.05, most *P* < 0.001; Table 2).

### Genetic architecture of plant and caterpillar traits

The *M. truncatula* SNP data explained a modest to substantial proportion of trait variation (Table S2, Fig. 2). On average, *M. truncatula* genetic variation accounted for a greater proportion of the variation in plant morphology traits (mean PVE = 0.40) than in IR traits (mean PVE = 0.17) or caterpillar performance (mean PVE = 0.24; recall that PVE is similar to narrow-sense heritability). However, *M. truncatula* genetics explained a particularly large amount of the variation in *L. melissa* caterpillar 16-day weight (PVE = 0.41, 90% equal-tail probability intervals [ETPIs] = 0.34–0.49; this trait also exhibited high among-line variance, Table 2). Estimates of PVE were generally precise, such that the average width of the 90% ETPIs for these parameters (mean across traits) was 0.13 (range = 0.11–0.15). In contrast, our estimates of the number of genetic loci with measurable effects on each trait (n-γ), and of the proportion of the PVE explained by those loci (PGE) were less certain; in particular, the average width of the 90% ETPIs for n-γ and PGE (a proportion) were 153.7 loci and 0.82, respectively (Table S2). Thus, uncertainty in these parameter estimates blurs differences in genetic architectures among traits suggested by the differences in parameter point estimates (compare Fig. 2 with Table S2). Genetic architecture parameter estimates for permuted (randomized) caterpillar performance data differed markedly from those for the actual data, most notably in terms of PVE. Whereas permutations of the survival to eclosion data did sometimes give modest estimates of PVE (the maximum was 0.12, 90% ETPIs = 0.06–0.19), these were still lower than the PVE estimate for the least heritable trait, namely survival to eclosion (PVE = 0.15, 90% ETPIs = 0.09–0.22), and most PVE estimates from permuted data were less than 0.05 (Fig. S4).

Consistent with the high (but uncertain) estimates of n-γ for most traits, many SNPs had small but non-zero posterior inclusion probabilities (PIPs) in the BSLMMs (Fig. S5). In other words, we were better able to detect than confidently isolate and localize the effects of individual genetic loci on the traits. There were a few exceptions to this pattern, most notably plant height and survival to eclosion. For plant height, one SNP each on chromosomes 5 and 7 had very high PIPs, ~ 1.0 (Fig. S6). Two nearby SNPs on chromosome 6 were confidently associated with survival to eclosion, but given the unbalanced design (most caterpillars did not survive to eclosion) and the modest difference between PGE (and to a lesser extent PVE) estimates for this trait and permutations of this trait, we do not interpret or discuss these associations with survival further. We next summarized the genomic distribution of genetic variants affecting each trait by estimating the number of QTL (or QTN) for each trait on each of the eight *M. truncatula* chromosomes (as in Santure *et al.*, 2015; Lucas *et al.*, 2018). This was done by summing the PIPs across all SNPs on each chromosome, and thus is analogous to the parameter n-γ, except that it is refers to specific chromosomes rather than the whole genome (Guan & Stephens, 2011; Riesch *et al.*, 2017; Lucas *et al.*, 2018). As these chromosomes vary little in size (~35 to 55 megabases), the number of QTL per chromosome should be similar across chromosomes if the traits are highly polygenic. Consistent with this prediction, evidence of putative QTL for most traits was not restricted to specific chromosomes but distributed relatively evenly among chromosomes (Figs. 3, S7).

**Figure 3:**
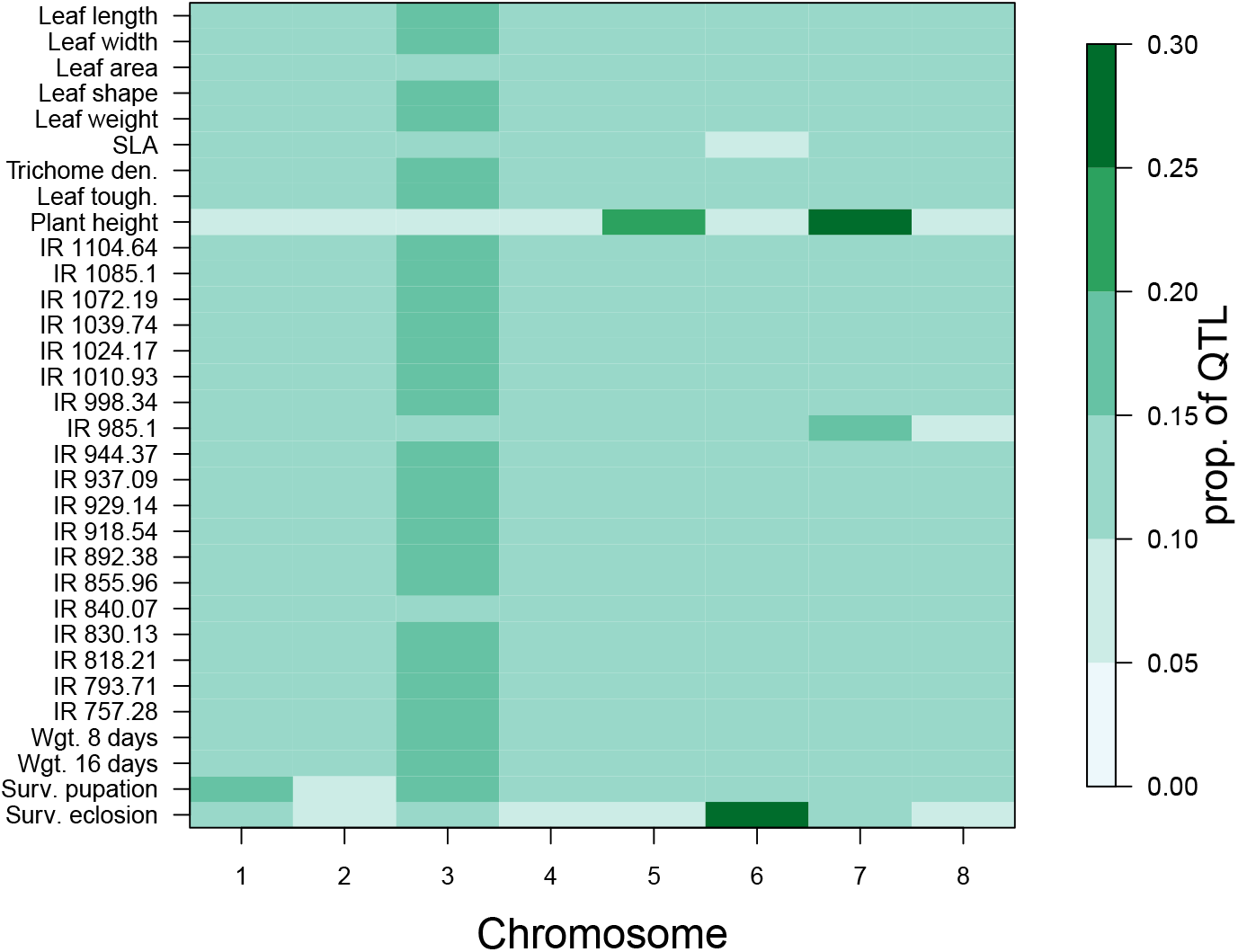
Heatmap image showing the proportion of QTL estimated for each trait on each of the eight *M. truncatula* chromosomes. The number of QTL per chromosome was estimated as the sum of the posterior inclusions probabilities across all SNPs on each chromosome. This was then divided by the total (sum) across chromosomes to obtain the proportions. For most traits, the genetic signal (i.e., QTL) were spread uniformly across chromosomes (also see Fig. S5), but for a few traits, especially plant height and survival to eclosion, QTL were clustered on one or a few chromosomes (also see Fig. S6). Note that chromosome 3 is slightly larger than the other chromosomes and thus harbors a slight excess of QTL for most traits. See Fig. S7 for numbers of QTL on each chromosome.

### Relationship between plant trait genetics and caterpillar performance

Trait genetic covariances and correlations were high for some pairs or sets of traits (high genetic correlations imply pleiotropy or tight linkage of causal variants; Fig. 4). For example, genetic correlations among leaf length, width, area and dry weight were all *r* ≥0.8. High, positive genetic correlations were also observed among the caterpillar performance traits, particularly 16-day weight, survival to pupation and survival to eclosion (*r* = 0.47 to 0.60). Caterpillar performance traits also exhibited non-trivial genetic correlations with several plant traits, most notably with leaf toughness where genetic correlations ranged from −0.25 for 8 day weight (95% confidence intervals [CIs] = −0.43 to −0.05, *P* = 0.016) to −0.39 for 16 day weight (95% CIs = −0.55 to −0.21, *P* < 0.001; Fig. 4). Weaker, but still consistently negative genetic correlations were observed between caterpillar performance traits and both trichome density and plant height (Fig. S8). More generally, hierarchical clustering revealed sets or modules of traits with high (positive or negative) genetic correlations, particularly for suites of IR spectra traits (Fig. S9).

**Figure 4:**
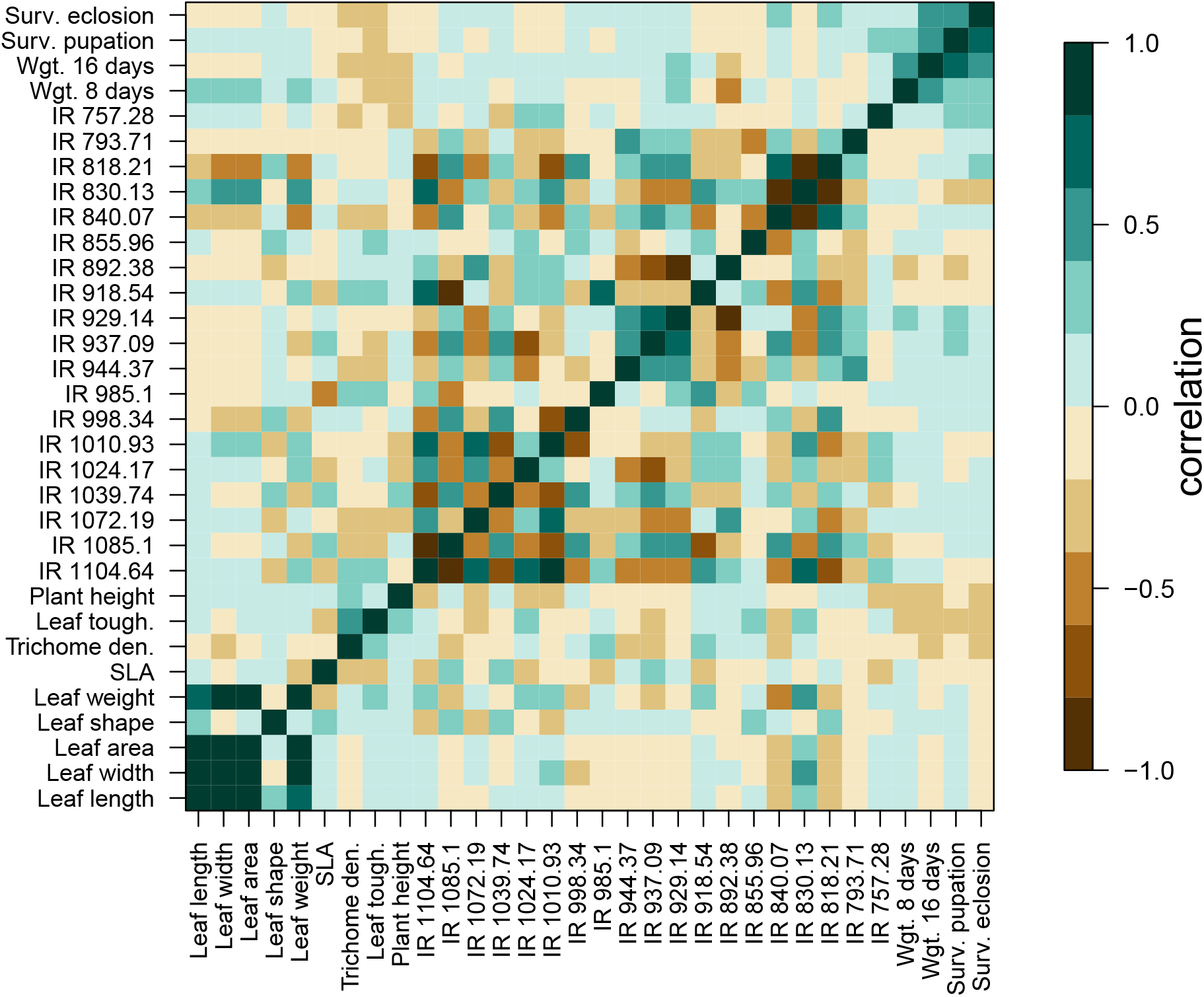
Heat map of (additive) genetic (upper triangle) and mean phenotype (lower triangle) Pearson correlation coefficients for pairs of plant and caterpillar traits. Genetic correlations were computed from genomic estimated breeding values (GEBVs) and mean phenotype correlations were computed using the phenotypic means of each trait for each plant line. Genetic and mean phenotype correlations were highly correlated with one another (*r* = 0.98).

Multiple regression models with Bayesian model averaging had some (albeit modest) predictive power, with correlations between observed and predicted caterpillar performance GEBVs ranging from *r* = 0.12 for survival to pupation (i.e., *r*^2^ = 1.4% of the variation in observed GEBVs explained by predictions) to r = 0.42 for 8-day weight (*r*^2^ = 17.6% of the variation in the observed GEBVs explained by predictions; Fig. 5). The most important predictor for 8-day caterpillar weight was IR 892.38, followed by IR 1072.19 (IR traits are labeled by their wavelength in cm^−1^; Figs. 5, S10). In contrast, leaf toughness was the best predictor of the GEBVs for 16-day weight, survival to pupation and survival to eclosion; higher GEBVs for leaf toughness consistently and credibly predicted lower GEBVs for caterpillar performance metrics. Leaf toughness was the only credible predictor of caterpillar survival (all other traits had PIPs < 0.5), whereas leaf toughness and several IR traits (or more precisely the GEBVs for these traits) had credible effects on 16 day weight GEBVS (i.e., IR 998.34, IR 1104.64 and IR 892.38; Figs. 5, S10). Standard multiple regression models that included the most credible covariates (those with PIP > 0.5; Fig. S10) explained 40.4% (8-day weight; covariates = IR 892.38 and IR 1072.19), 34.9% (16-day weight; covariates = leaf toughness, IR 998.34, IR 1004.64 and IR 892.38), 8.5% (survival to pupation; covariate = leaf toughness) and 12.1% (survival to eclosion; covariate = leaf toughness) of the variation in caterpillar performance GEBVs, with all included covariates having significant effects (all *P* < 0.01). Thus, models with the most important covariates explained a moderate amount of the variation in caterpillar performance GEBVs, but still less than 50% in all cases.

**Figure 5:**
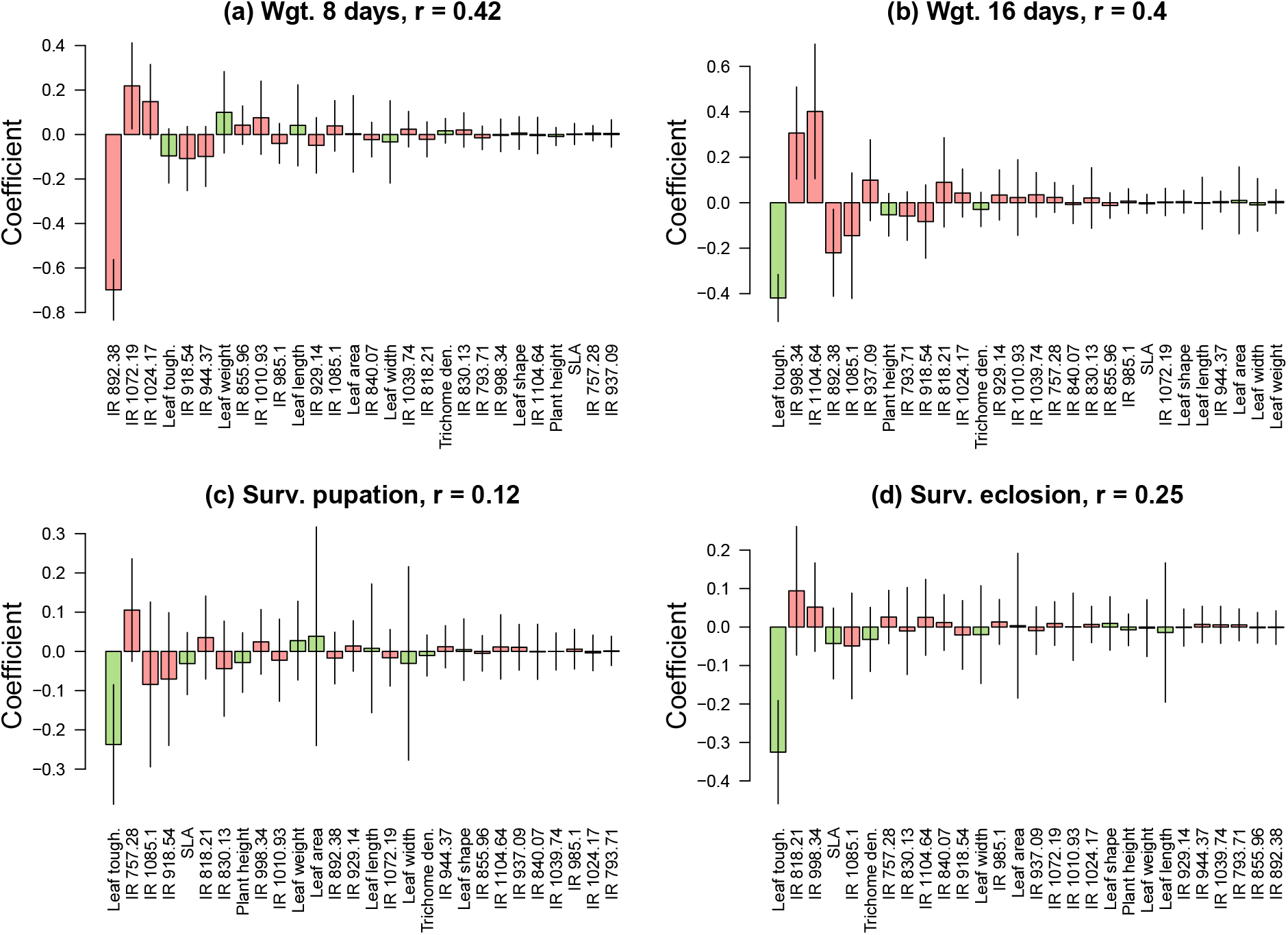
Barplots showing the effect of the genetic component of plant traits on the genetic component of caterpillar performance, specifically (a) weight at 8 days, (b) weight at 16 days, (c) survival to pupation, and (d) survival to eclosion. Bars denote Bayesian model-averaged estimates (posterior means) of standardized regression coefficients for the effect of the genomic estimated breeding values (GEBVs) for each plant trait on the GEBVs for the caterpillar performance traits. Traits are sorted by the absolute magnitude of these estimates. Vertical bars denote ± one standard deviation of the posterior (analogous to a standard error). Colors distinguish between plant growth and defense traits (green) and IR traits (pink). Pearson correlations between the caterpillar performance GEBVs and estimates of these from 10-fold cross-validation are given in the panel headers (see the main text for corresponding *r*^2^ values). See Fig. S10 for covariate posterior inclusion probabilities.

For 8-day caterpillar weight GEBVs, predictions from random forest regression accounted for 31.9% of out-of-bag (OOB) variance (OOB variance measures predictive performance) (`mtry` = 18, `nodesize` = 2). The most important predictor variables were IR 892.38, IR 985.1, and plant height (Fig. 6a). For 16-day caterpillar weight GEBVs, random forest explained 14.4% of the OOB variance (`mtry` = 12, `nodesize` = 9). The most important predictor variables in this case were leaf toughness, IR 1104.64, and IR 830.13 (Fig. 6b). Only 5.3% of the OOB variance was explained for survival to eclosion, with leaf toughness and IR 830.13 being the most important traits (Fig. 6c). Graphical analyses of the random forest regression results suggested non-linear relationships between GEBVs for many of the top plant and caterpillar traits (Figs. 6d-f, S11 and S12). For example, the effects of IR 892.38 and IR 985.1 on 8-day weight exhibited a strong interaction (a similar pattern held for many of the IR chemical features). In contrast, the effect of leaf toughness on 16-day weight was negative and nearly linear (tougher leaves were associated with lower weights), although there was evidence of an asymptote at higher values of leaf toughness. We failed to explain a non-zero proportion of the OOB variance in caterpillar survival to pupation with random forest regression, and thus results for this trait are not shown.

**Figure 6:**
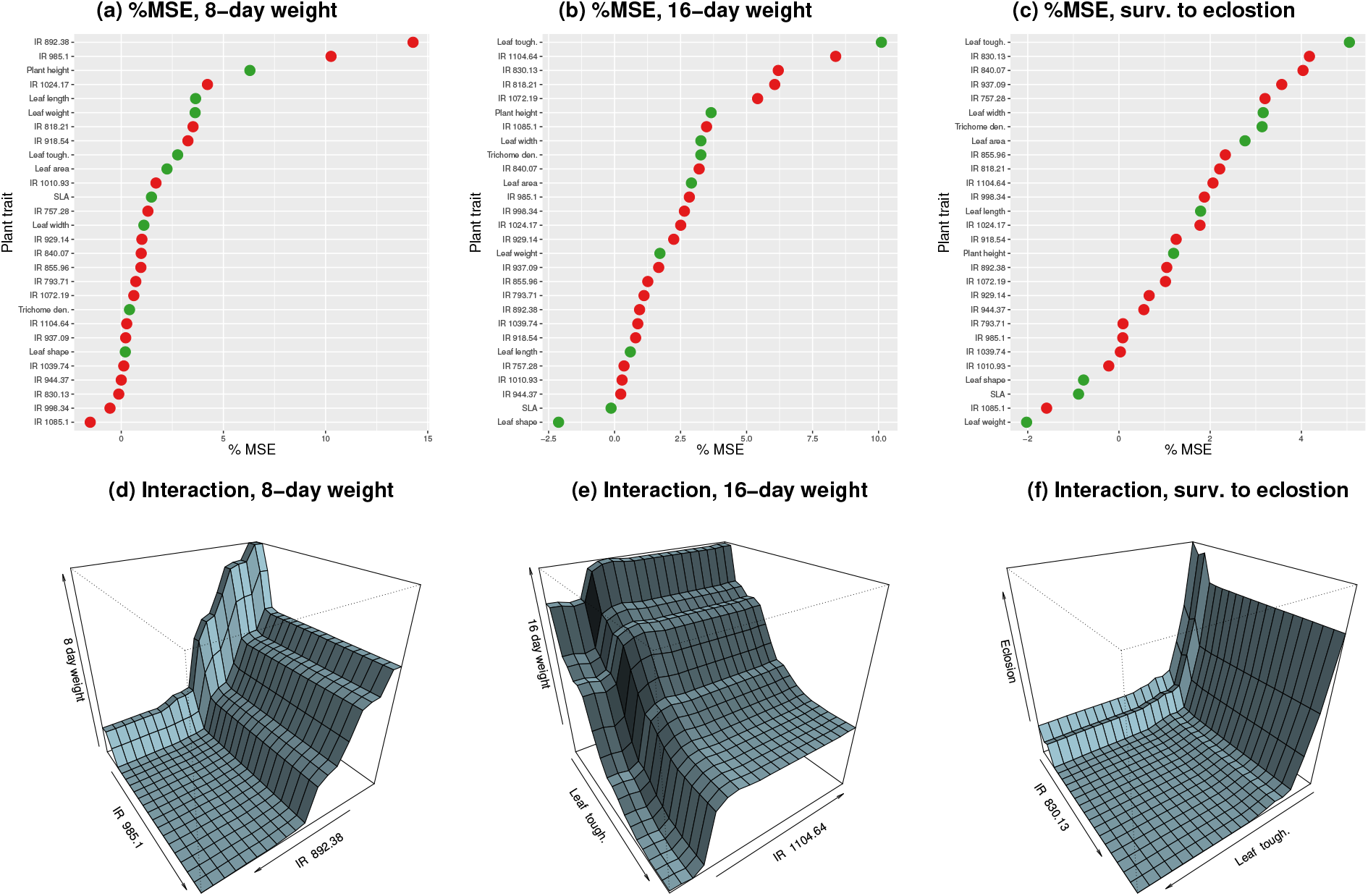
Summary of random forest analysis for predicting caterpillar GEBVs from plant trait GEBVs. Panels (a-c) show the importance of each covariate (plant trait GEBV), and panels (d-f) depict the relationships between the two most important covariates and GEBVs for caterpillar weight at 8 (d) or 16 (e) days and survival to eclosion (f). Plots in d-f were completed with `plotmo` (Milborrow, 2018) and illustrate interactions between the top two predictor variables. See Figs. S11 and S12 for additional interactions.

## Discussion

Because the world is full of newly-formed host-parasite interactions (including plant-insect interactions involving consumputive herbvory; Nylin *et al.*, 2018), and because most novel host plants are relatively sub-optimal hosts (Yoon & Read, 2016), the results reported here are of interest not only as a step towards understanding the interaction between *L. melissa* and *M. sativa* (discussed further below), but also as a more general model for the formation of host-parasite interactions. In addition, genetic dissections of plant-insect interactions are important not only for understanding the complexity underlying the formation and persistence of new associations, but also for understanding the evolution of plant defensive traits and phytochemical diversity in terrestrial ecosytems. In our study, genetic variation within *M. truncatula* explained a non-trivial proportion of the variation in *L. melissa* caterpillar performance traits, especially 16-day weight (PVE = 0.41) and survival to pupation (PVE = 0.31). Estimates of the variance in plant and caterpillar traits explained (PVE) by plant genetic variation were similar, meaning the two sets of traits were (on average) similarly heritable with respect to *M. truncatula* (this suggests these caterpillar performance traits can meaningfully be viewed as extended phenotypes of *M. truncatula, sensu* Dawkins, 1982; see also, e.g., Whitham *et al.*, 2006).

Genomic estimated breeding values (GEBVs) for caterpillar performance traits were most consistently and strongly associated with GEBVs for leaf toughness, with more modest or idiosyncratic correlations with several IR chemical features (e.g., IR 892.38 and IR 1104.64), trichome density, and plant height. These genetic correlations suggest either that caterpillar performance and several of these plant traits are affected by some of the same segregating genetic variants (i.e., pleiotropy), or that modest to high LD exists among genetic variants affecting the plant traits and caterpillar performance. Such high LD would imply tight linkage among many genetic variants, or some alternative process or mechanism for suppressed recombination among genotypes (this could include low rates of gene flow among the natural source populations from which these lines were derived). However, LD is modest and decays with a few kbs to background levels in this mapping population (i.e., mean LD, measured by *r*^2^ drops below 0.2 within 20 kbs; Branca *et al.*, 2011). Interestingly, the additive effects of alleles on the measured plant traits (as captured by the trait GEBVs) were able to explain or account for the additive effects of *M. truncatula* alleles on caterpillar performance, at least to a modest extent (as expected, explanatory power was lower for cross-validation than in simple linear models). Nonetheless, much of the variation in caterpillar performance GEBVs was not accounted for by the plant trait GEBVs. This implies additional plant traits (and underlying genes) likely contribute to the total variation in caterpillar performance explained by plant genetics. We discuss these results in more detail below.

### The genetic architecture of traits associated with a plant-insect interaction

Our results were consistent with standing genetic variation at many loci in *M. truncatula* for *L. melissa* caterpillar performance on *M. truncatula*. Specifically, estimates of PVE from the BSLMMs and REML estimates of the among plant-line genetic variances provide direct evidence of standing polygenic variation in *M. truncatula* for *L. melissa* caterpillar performance. Furthermore, results from the BSLMMs suggest multiple QTL for caterpillar performance are dispersed across the eight *M. truncatula* chromosomes rather than localized in one or a few regions of the genome. A polygenic basis for caterpillar performance (as a plant trait) was also detected in a recent genomic study of *Pieris rapae* caterpillars reared on *Arabidopsis thaliana* (Nallu *et al.*, 2018). In this study, Nallu *et al.* (2018) identified 12 *A. thaliana* genes associated with variation in *P. rapae* performance (weight gain over 72 hours), which included CYP79B2, a cytochrome P450 gene known to affect plant resistance to insects. A genome-wide transcriptomic response to herbivory (and even to oviposition) was detected as well.

More generally, genetic variation for resistance to insects has been documented in numerous other plant species, especially crops (Via, 1990; Schoonhoven *et al.*, 2010), although mostly without genome-scale data and without explicit links to plant traits. Still, these studies show that intraspecific variation in plant resistance to insects is often highly heritable, and that it can involve one or many genes (reviewed in Schoonhoven *et al.*, 2010). The same plant species can even exhibit polygenic resistance variation with respect to one insect species and monogenic resistance variation with respect to another (Kennedy & Barbour, 1992). Thus, while our finding of a polygenic architecture is not unexpected given the complex, multifaceted nature of caterpillar performance (Allen *et al.*, 2010; Rockman, 2012), additional genomic studies are needed for a more robust assessment of the prevalence and consistency of this pattern (especially in natural systems).

The full set of plant and caterpillar traits we measured exhibited a range of heritabilities, yet, with the possible exception of plant height, we found little evidence of major effect loci. Instead the traits appeared to be controlled by many loci. Genome-wide association mapping methods (and to a lesser extent genomic prediction methods) are known to suffer from a failure to detect many small effect variants (Eichler *et al.*, 2010; Yang *et al.*, 2010), and from overestimating the effects of large effect variants (i.e., the Beavis effect; Beavis, 1998). However, major-effect loci are less likely to be missed. This is true in general as such loci are easier to detect even with small sample sizes, but especially true here given the high-density genome-wide SNP data set we used (>five million SNPs, or about one per 100 bps) and thus the high likelihood of LD between at least one of our SNPs and most causal variants. Moreover, two of the plant traits we analyzed, plant height and trichome density, were independently mapped and analyzed in an earlier study of the *M. truncatula* HapMap mapping population (albeit with a different subset of lines) (Stanton-Geddes *et al.*, 2013). Results from Stanton-Geddes *et al.* (2013) and our results were remarkably consistent, with, for example, 58% versus 59% (plant height) and 45% versus 49% (trichome density) of the trait variation partitioned among lines in Stanton-Geddes *et al.* (2013) versus our study, respectively. This is reassuring, particularly given the variability frequently observed in genetic mapping and quantitative genetic results among mapping populations and environments (e.g., Weinig *et al.*, 2002, 2003; Weiss, 2008). However, the use of inbred lines sampled from many localities necessarily distorts the frequencies and possibly average effects of genetic variants on traits, thus our results do not rule out major-effect loci for these traits in natural populations.

### Evidence of pleiotropic effects across species, and of variance left unexplained

The estimated genetic correlations are consistent with either pleiotropic effects of *M. truncatula* alleles on plant traits and caterpillar performance, or with LD among variants that independently affect subsets of these traits (parsing these two possibilities is very difficult, and at the extreme, very tight linkage can be functionally equivalent to pleiotropy). Leaf toughness, and to a lesser extent, trichome density and plant height, exhibited some of the greatest and most consistent negative genetic correlations with *L. melissa* performance. Leaf toughness and trichome density constitute structural (physical) plant defenses (Levin, 1973; Schoonhoven *et al.*, 2010), and our results thus support recent calls for greater attention to structural (as opposed to chemical) plant defenses (Hanley *et al.*, 2007; Carmona *et al.*, 2011; Malishev & Sanson, 2015). However, some IR chemical features exhibited high genetic correlations with some or many of the caterpillar performance traits. This is consistent with a role for intraspecific variation in phytochemical defenses in *M. truncatula* as well, although the IR chemical features could also reflect variation in plant nutritional composition rather than chemical defenses *per se.* Future work should identify the molecules underlying variation at the leading IR chemical features (e.g., IR 892.38 and IR 1104.64).

Plant trait GEBVs accounted for a moderate amount of the variation in caterpillar weight GEBVs, but relatively little of the variation in caterpillar survival GEBVs. In other words, our results suggest that the alleles affecting the measured plant traits accounted for a greater proportion of the heritable variation in *M. truncatula* for caterpillar weight than caterpillar survival. Nonetheless, in no cases did the variance explained or predictive power of these models approach 100%. In fact, the highest percent variance explained was 40.8%, and predictive power never exceeded 17.6% for the Bayesian multiple regression or 31.9% for the random forest regression. This means that the effects of *M. truncatula* alleles on caterpillar performance are not fully accounted for by the effects of these alleles on the measured plant traits. Additional heritable plant traits not measured in this study must affect *L. melissa* performance, and additional work will be required to identify these. Obvious candidates include defensive phytochemicals or plant nutrients that were not captured by the IR assays. Still, even the modest predictive power of these models allows us to conclude, for example, that the genetic quality of a plant in terms of caterpillar performance can be predicted in part from the additive effects of plant alleles on leaf toughness.

As expected, the plant traits most important in these predictive models tended to be the ones with the largest genetic correlations with caterpillar performance. However, there were a few exceptions that arose because of correlations among the plant trait GEBVs, which rendered a subset of these traits (e.g., trichome density) unimportant in the predictive models. Moreover, the relative ranks of plant traits in terms of their importance (i.e., Bayesian model-averaged effect estimates or percent reduction in MSE) differed between the Bayesian multiple regression models and random forest regression. We think these differences were most evident in cases where random forest regression identified extreme interactions among plant trait GEBVs or non-linear relationships between GEBVs for the plant traits and caterpillar performance (e.g., IR 985.1 on 8-day caterpillar weight), as these would not be captured by the Bayesian multiple regression models.

### Conclusions and future directions

We have shown that plant genetic variation can have a substantial effect on the outcome of a plant-insect interaction, specifically on whether *L. melissa* caterpillars can develop successfully on *M. truncatula*. Genetic variation among *M. truncatula* plants explained about as much of the variance in caterpillar performance in the current study (9-41%) as genetic variation among *L. melissa* caterpillars did in an earlier rearing experiment on *M. sativa* (7-57%) (Gompert *et al.*, 2015). This suggests that caterpillar and plant genetic variation combined could explain a large proportion (i.e., over half) of the variation in larval performance, which is necessarily a key aspect of the interaction between plants and herbivorous insects. However, *M. truncatula* and *M. sativa* are not identical, and it remains to be seen whether similar levels of genetic variation for performance exist in this actual (rather than potential) *L. melissa* host plant. Moreover, gene by gene epistatic interactions between *L. melissa* alleles and *M. sativa* (or *M. truncatula*) alleles could modulate the total variance in performance explained for the pair of species (in other words, the trait heritabilities with respect to plant and insect genes are not necessarily additive).

Ultimately, we want to accurately predict the mosaic patterns of host use and host adaption in *L. melissa* from a mechanistic understanding of the factors affecting host use. We have reasons to be both optimistic and pessimistic about this aim. Past work on *L. melissa* has shown that genetic variants associated with performance in the lab covary significantly with host use in nature (Gompert *et al.*, 2015; Chaturvedi *et al.*, 2018). Thus, genetic variants affecting performance in the lab appear to also be associated with host-plant adaptation in nature. On the other hand, the lab environment is necessarily simplified and lacks interactions with predators, competitors and mutualists that could be important determinants of host use in the wild. For example, survival of *L. melissa* caterpillars on *M. sativa* in a field experiment depended on the presence of ants that defend the caterpillars from predators (this is a facultative relationship where the ants receive a sugar reward from the caterpillars; Forister *et al.*, 2011). Even ignoring such complexities, the relevance of genetic and trait variation in *M. truncatula* for understanding genetic and trait variation in *M. sativa* is not certain. Leaf toughness, which was most strongly associated with performance in the current experiment, exhibits a similar range of variation in *M. sativa* and *M. truncatula* (albeit with somewhat tougher leaves in *M. sativa* on average; Harrison *et al.*, 2018). This suggests variation in leaf toughness in *M. sativa* could have a similar affect on caterpillar performance. In the end, we may fail to generate reliable predictions about host use in nature from simple lab experiments, but nonetheless might advance scientific understanding of the importance of intraspecific variation for the evolution and ecology of plant-insect interactions by gaining a better understanding of how and why these predictions fail.

## Acknowledgments

We thank Shaylen Fidel, Camden Treat, Marissa Kee and Kenneth Tache for their help with the caterpillar rearing experiment, and Marianne Harris for use of greenhouse space and help caring for the plants. Chris Nice, James Fordyce, Alex Buerkle and Amy Springer provided valuable comments on an earlier draft of this manuscript. This research was funded by the National Science Foundation (DEB-1638768 to ZG and DEB-1638793 to MLF), and by the Utah Agricultural Experiment Station (paper no. XXX). The support and resources from the Center for High Performance Computing at the University of Utah are also gratefully acknowledged.

## Data Accessibility

All original data and scripts will be deposited on Dryad.

## Author Contributions

ZG, LKL and MB designed the study. ZG, LKL, MB, FZ, CP, MJT and MLF conducted the experiment. ZG, FC, CP and TS analyzed the data. ZG, CP and TS wrote the manuscript. All authors revised and edited the manuscript.

## Supplemental material for

### Supplemental Methods and Results

#### Planting and tending *Medicago truncatula*

Our methods for planting and growing *Medicago truncatula* were developed based on https://www.noble.org/globalassets/docs/medicago-handbook/growing-medicago-truncatula.pdp and https://www.noble.org/globalassets/docs/medicago-handbook/vernalization.pdf. As described in the main text, we first mechanically scarified the seeds with sandpaper, and then placed five seeds from each plant line in 4in × 4in × 3.5in pots with a 4:1 mixture of Sunshine Mix #4 soil and Perlite. Seeds were placed on top of wet soil in a slight divot, and then covered with ~10mm mixture of a dry soil and Perlite. We then misted the pots and covered them with humidity domes until germination. 10 pots were placed under each humidity dome, and the placement of pots (within and among domes amnd trays) was randomized within each replicate (i.e., block). Plants were thinned on May 26th (i.e., after germination was complete) to ensure that no pots had more than two plants. This was done to minimize competition among plants, while still providing sufficient plant biomass for our caterpillar rearing experiments. Ladybugs were introduced into the greenhouse on July 8th and 9th for biological control of aphids and other pests.

#### BSLMMs fit to randomized data

Past work has shown that BSLMMs provide a robust method for genome-wide association mapping and genomic prediction (e.g., Zhou *et al.*, 2013; Gompert *et al.*, 2017), even when modeling binary traits (Guan & Stephens, 2011). However, we were concerned that the models might perform poorly when presented with binary data where most individuals had either 0s or 1s, as is the case for our survival data (particularly survival to eclosion). To assess this possibility, and specifically to verify that our results would not be expected for random phenotypic data, we analyzed 12 pseudo-data sets. We obtained these 12 data sets by randomizing the trait data for each of the four caterpillar performance traits three times (generating 12 randomized data sets total). Thus, half of these data sets were based on the binary survival data, and half on more standard quantitative data. We then fit BSLMMs for each of the 12 pseudo-data sets using `gemma` (version 0.94.1; Zhou *et al.*, 2013) with 15 MCMC runs each with a 500,000 iteration burn-in followed by 2 million sampling steps with a thinning interval of 20. As with the actual data sets, we only considered SNPs with a minor allele frequency greater than 0.01. Results from these analyses are shown in Fig. S4 and presented in the main text of this manuscript.

#### Supplemental Tables and Figures

**Table S1:** Hapmap IDs, population codes, countries of origin, and seed sources for the 94 *Medicago truncatula* lines used in this study. Additional information about these inbred lines and the *M. truncatula* Hapmap project is available from http://www.medicagohapmap.org/home/view.

| ID | Population | Country | Source |
| --- | --- | --- | --- |
| HM001 | SA22322 | Syria | INRA-Montpellier |
| HM003 | ESP105-L | Spain | INRA-Montpellier |
| HM004 | DZA045-6 | Algeria | INRA-Montpellier |
| HM008 | DZA012-J | Algeria | INRA-Montpellier |
| HM009 | GRC020-B | Greece | INRA-Montpellier |
| HM010 | SA24714 | Italy | INRA-Montpellier |
| HM011 | DZA327-7 | Algeria | INRA-Montpellier |
| HM012 | SA26063 | Morocco | INRA-Montpellier |
| HM027 | F83005-9 | France | INRA-Montpellier |
| HM032 | F11005-E | France | INRA-Montpellier |
| HM035 | F66017 | France | INRA-Montpellier |
| HM039 | SA03116 | Israel | INRA-Montpellier |
| HM040 | SA03780 | Italy | INRA-Montpellier |
| HM041 | SA09048 | Libya | INRA-Montpellier |
| HM044 | SA14161 | Jordan | INRA-Montpellier |
| HM046 | SA27882 | Morocco | INRA-Montpellier |
| HM048 | DZA016-F | Algeria | INRA-Montpellier |
| HM049 | DZA058-5 | Algeria | INRA-Montpellier |
| HM055 | DZA326 | Algeria | INRA-Montpellier |
| HM058 | ESP163-E | Spain | INRA-Montpellier |
| HM060 | F20015-10 | France | INRA-Montpellier |
| HM061 | GRC033-B2 | Greece | INRA-Montpellier |
| HM065 | PRT179-J | Portugal | INRA-Montpellier |
| HM070 | SA08625 | Morocco | INRA-Montpellier |
| HM073 | SA09710 | Tunisia | INRA-Montpellier |
| HM076 | SA23859 | Tunisia | INRA-Montpellier |
| HM079 | DZA045-4c | Algeria | INRA-Montpellier |
| HM080 | DZA061-B3d | Algeria | INRA-Montpellier |
| HM081 | DZA202-5 | Algeria | INRA-Montpellier |
| HM087 | DZA323-1 | Algeria | INRA-Montpellier |
| HM091 | ESP171-F | Spain | INRA-Montpellier |
| HM105 | SA09137 | Algeria | INRA-Montpellier |
| HM106 | SA09434 | Tunisia | INRA-Montpellier |
| HM108 | SA09715 | Tunisia | INRA-Montpellier |
| HM111 | SA27192 | Italy | INRA-Montpellier |
| HM115 | Cyprus_C | Cyprus | INRA-Montpellier |
| HM117 | ESP031-A | Spain | INRA-Montpellier |
| HM120 | ESP095-C | Spain | INRA-Montpellier |
| HM127 | F20025-F | France | INRA-Montpellier |
| HM130 | F20069-C | France | INRA-Montpellier |
| HM135 | SA02748 | Israel | INRA-Montpellier |
| HM139 | SA08623 | Morocco | INRA-Montpellier |
| HM143 | SA10481 | Tunisia | INRA-Montpellier |
| HM146 | SA21302 | Libya | INRA-Montpellier |
| HM150 | SA22323 | Syria | INRA-Montpellier |
| HM163 | DZA061-11 | Algeria | INRA-Montpellier |
| HM165 | DZA231-1 | Algeria | INRA-Montpellier |
| HM177 | ESP100-G | Spain | INRA-Montpellier |
| HM179 | ESP162-A | Spain | INRA-Montpellier |
| HM180 | ESP163-C | Spain | INRA-Montpellier |
| HM184 | F20058-B | France | INRA-Montpellier |
| HM186 | GRC024-H | Greece | INRA-Montpellier |
| HM194 | SA09700 | Tunisia | INRA-Montpellier |
| HM195 | SA09728 | Tunisia | INRA-Montpellier |
| HM196 | SA09970 | Tunisia | INRA-Montpellier |
| HM199 | SA19983 | Cyprus | INRA-Montpellier |
| HM202 | SA25941 | Italy | INRA-Montpellier |
| HM205 | SA28375 | Portugal | INRA-Montpellier |
| HM253 | PI660496SSD | France | USDA-ARS |
| HM256 | PI442895SSD | Australia | USDA-ARS |
| HM259 | PI577599SSD | Greece | USDA-ARS |
| HM260 | PI516934SSD | Morocco | USDA-ARS |
| HM262 | PI564941SSD | Morocco | USDA-ARS |
| HM266 | PI660450SSD | Algeria | USDA-ARS |
| HM267 | PI660437SSD | Algeria | USDA-ARS |
| HM268 | PI660438SSD | Algeria | USDA-ARS |
| HM269 | PI660470SSD | Unknown | USDA-ARS |
| HM270 | PI493297SSD | Portugal | USDA-ARS |
| HM271 | PI384664SSD | Morocco | USDA-ARS |
| HM276 | PI577627SSD | Algeria | USDA-ARS |
| HM277 | PI577621SSD | Algeria | USDA-ARS |
| HM279 | PI660460SSD | Morocco | USDA-ARS |
| HM287 | PI577607SSD | Lebanon | USDA-ARS |
| HM288 | PI577611SSD | Germany | USDA-ARS |
| HM289 | PI577617SSD | Greece | USDA-ARS |
| HM290 | PI577640SSD | U.S. | USDA-ARS |
| HM293 | PI660411SSD | Italy | USDA-ARS |
| HM294 | PI660433SSD | Algeria | USDA-ARS |
| HM295 | PI660442SSD | Algeria | USDA-ARS |
| HM296 | PI660444SSD | Algeria | USDA-ARS |
| HM297 | PI660447SSD | Algeria | USDA-ARS |
| HM298 | PI660448SSD | Algeria | USDA-ARS |
| HM299 | PI660456SSD | Morocco | USDA-ARS |
| HM301 | PI660494SSD | Italy | USDA-ARS |
| HM302 | PI283662SSD | Italy | USDA-ARS |
| HM307 | PI516927SSD | Morocco | USDA-ARS |
| HM308 | PI516933SSD | Morocco | USDA-ARS |
| HM309 | PI516939SSD | Morocco | USDA-ARS |
| HM310 | PI535651SSD | Tunisia | USDA-ARS |
| HM311 | PI535752SSD | Morocco | USDA-ARS |
| HM312 | PI577609SSD | Sweden | USDA-ARS |
| HM314 | PI660361SSD | Greece | USDA-ARS |
| HM315 | PI660387SSD | France | USDA-ARS |
| HM316 | PI660421SSD | Australia | USDA-ARS |

**Figure S1:**
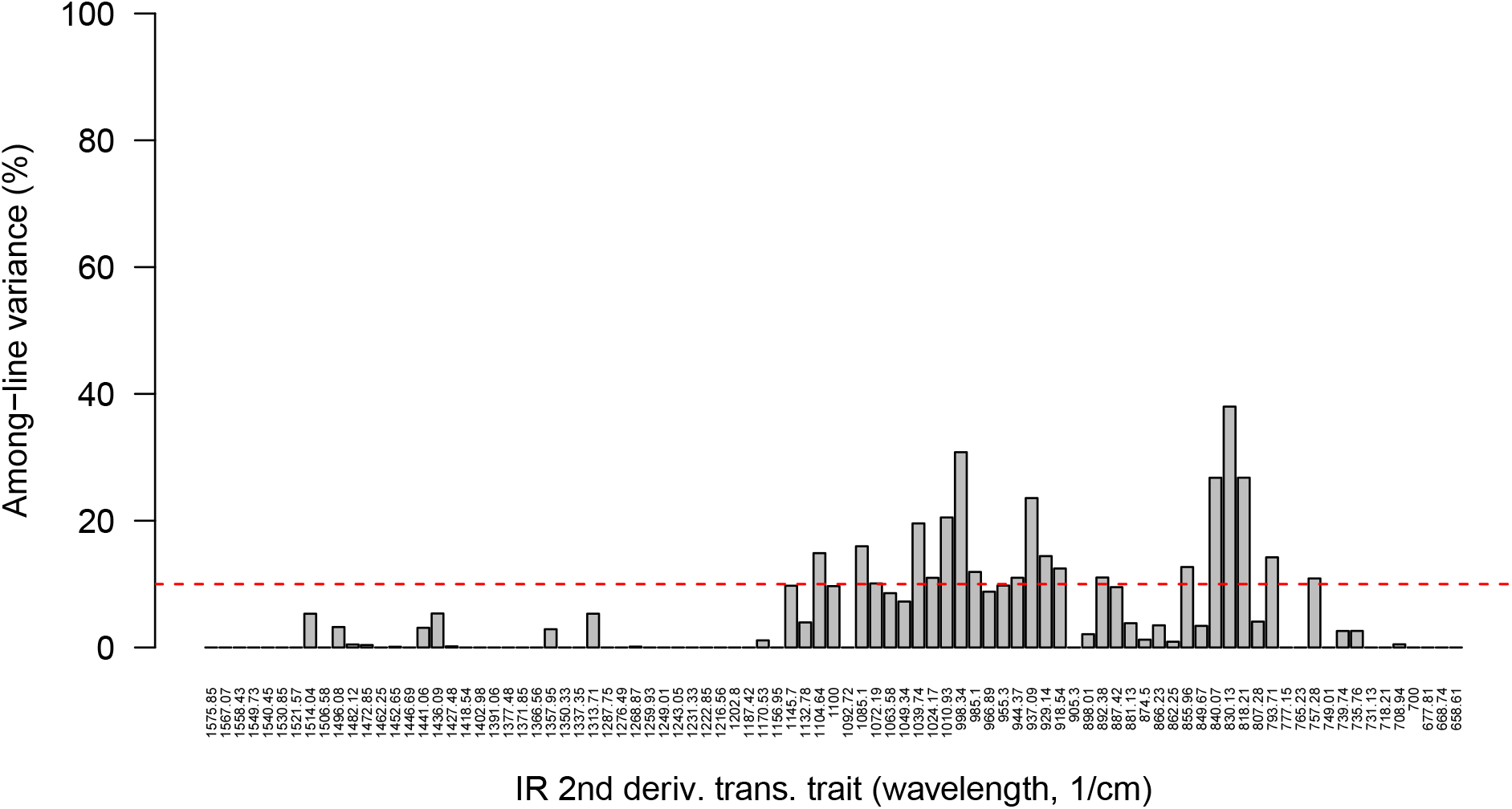
Barplot displaying the percent variance among *M. truncatula* lines (i.e., genotypes) for each infrared (IR) spectra trait. Here, the IR traits are the 2nd derivatives of the transmittance spectra. We only considered IR traits with ≥ 10% of the variance among lines for downstream analyses (denoted here with a horizontal dashed line).

**Table S2:** Summary of trait genetic architectures for each plant and caterpillar trait based on the posterior probability distributions from fitting Bayesian sparse linear mixed models. Parameters shown are the proportion of the trait variation explained by *M. truncatula* genetics (PVE), the proportion of the PVE attributable to loci with measurable effects (PGE), and the number of loci with measurable effects (no. QTL). Posterior distributions are summarized based on the median (Med.) and the lower bounds (5th %) and upper bounds (95th %) of the 90% equal-tail probability intervals (ETPIs).

| Trait | PVE |  |  | PGE |  |  | no. QTL |  |  |
| --- | --- | --- | --- | --- | --- | --- | --- | --- | --- |
|  | Med. | 5th % | 95th % | Med. | 5th % | 95th % | Med. | 5th % | 95th % |
| Leaf length | 0.43 | 0.36 | 0.50 | 0.31 | 0.00 | 0.91 | 35 | 1 | 257 |
| Leaf width | 0.48 | 0.41 | 0.54 | 0.43 | 0.01 | 0.94 | 28 | 1 | 204 |
| Leaf area | 0.48 | 0.41 | 0.55 | 0.43 | 0.01 | 0.92 | 18 | 1 | 174 |
| Leaf shape | 0.21 | 0.14 | 0.29 | 0.38 | 0.00 | 0.93 | 14 | 1 | 231 |
| Leaf weight | 0.40 | 0.33 | 0.47 | 0.24 | 0.00 | 0.88 | 24 | 0 | 177 |
| SLA | 0.16 | 0.09 | 0.23 | 0.83 | 0.30 | 0.99 | 4 | 1 | 14 |
| Trichome den. | 0.49 | 0.42 | 0.56 | 0.41 | 0.01 | 0.94 | 39 | 1 | 249 |
| Leaf tough. | 0.34 | 0.27 | 0.41 | 0.34 | 0.00 | 0.90 | 15 | 1 | 193 |
| Plant height | 0.57 | 0.51 | 0.63 | 0.76 | 0.62 | 0.93 | 5 | 2 | 14 |
| IR 1104.64 | 0.13 | 0.07 | 0.20 | 0.62 | 0.07 | 0.95 | 4 | 1 | 133 |
| IR 1085.1 | 0.14 | 0.08 | 0.21 | 0.63 | 0.21 | 0.96 | 3 | 1 | 20 |
| IR 1072.19 | 0.10 | 0.04 | 0.17 | 0.30 | 0.00 | 0.91 | 10 | 0 | 129 |
| IR 1039.74 | 0.15 | 0.08 | 0.23 | 0.41 | 0.00 | 0.93 | 23 | 1 | 200 |
| IR 1024.17 | 0.11 | 0.05 | 0.18 | 0.39 | 0.00 | 0.93 | 10 | 0 | 147 |
| IR 1010.93 | 0.17 | 0.10 | 0.24 | 0.35 | 0.00 | 0.92 | 29 | 0 | 247 |
| IR 998.34 | 0.29 | 0.22 | 0.36 | 0.39 | 0.01 | 0.93 | 30 | 1 | 237 |
| IR 985.1 | 0.14 | 0.09 | 0.20 | 0.80 | 0.52 | 0.98 | 2 | 1 | 6 |
| IR 944.37 | 0.11 | 0.05 | 0.18 | 0.38 | 0.00 | 0.94 | 15 | 0 | 211 |
| IR 937.09 | 0.23 | 0.17 | 0.31 | 0.32 | 0.00 | 0.91 | 25 | 0 | 212 |
| IR 929.14 | 0.16 | 0.10 | 0.23 | 0.41 | 0.00 | 0.94 | 11 | 1 | 135 |
| IR 918.54 | 0.12 | 0.06 | 0.18 | 0.65 | 0.15 | 0.96 | 3 | 1 | 123 |
| IR 892.38 | 0.14 | 0.07 | 0.20 | 0.32 | 0.00 | 0.92 | 13 | 0 | 147 |
| IR 855.96 | 0.14 | 0.07 | 0.20 | 0.28 | 0.00 | 0.90 | 18 | 0 | 194 |
| IR 840.07 | 0.23 | 0.16 | 0.30 | 0.40 | 0.00 | 0.93 | 14 | 1 | 207 |
| IR 830.13 | 0.35 | 0.28 | 0.42 | 0.81 | 0.28 | 0.98 | 7 | 2 | 267 |
| IR 818.21 | 0.25 | 0.18 | 0.32 | 0.61 | 0.03 | 0.96 | 15 | 2 | 195 |
| IR 793.71 | 0.15 | 0.09 | 0.22 | 0.37 | 0.00 | 0.93 | 24 | 1 | 243 |
| IR 757.28 | 0.08 | 0.02 | 0.15 | 0.54 | 0.01 | 0.95 | 4 | 1 | 96 |
| Wgt. 8 days | 0.09 | 0.05 | 0.15 | 0.49 | 0.00 | 0.97 | 10 | 0 | 67 |
| Wgt. 16 days | 0.41 | 0.34 | 0.49 | 0.27 | 0.00 | 0.90 | 25 | 1 | 202 |
| Surv. pupation | 0.31 | 0.25 | 0.37 | 0.99 | 0.93 | 1.00 | 5 | 3 | 8 |
| Surv. eclosion | 0.15 | 0.09 | 0.22 | 0.97 | 0.84 | 1.00 | 2 | 1 | 6 |

**Figure S2:**
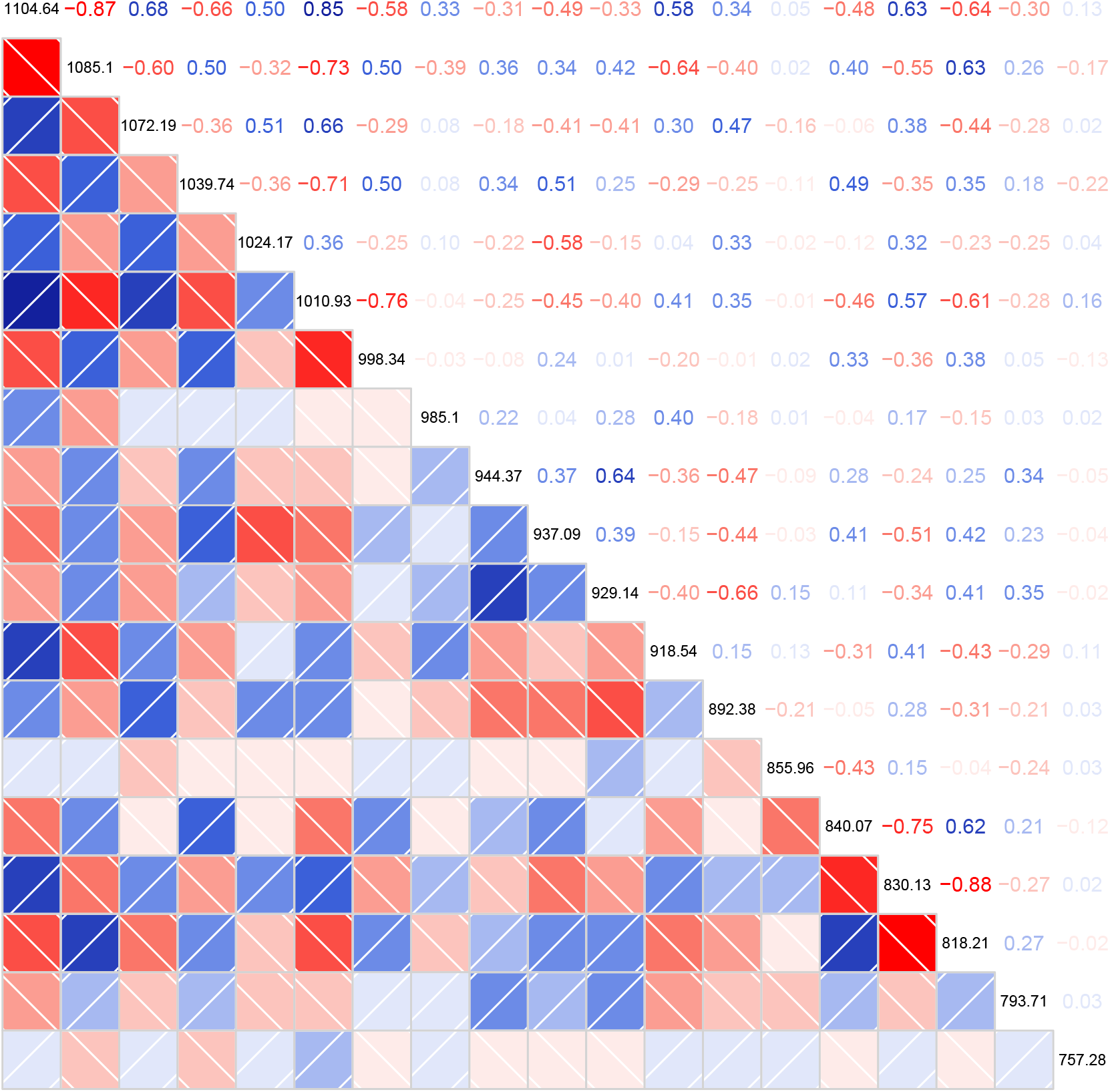
Correlogram giving pairwise phenotypic correlations for the 19 near-infrared spectra 2nd derivative traits. Pearson correlations are shown in the upper triangle of the correlation matrix, and depicted graphically in the lower triangle of the correlation matrix, with darker shading denoting higher correlations. Wavelengths defining each trait are given along the diagonal.

**Figure S3:**
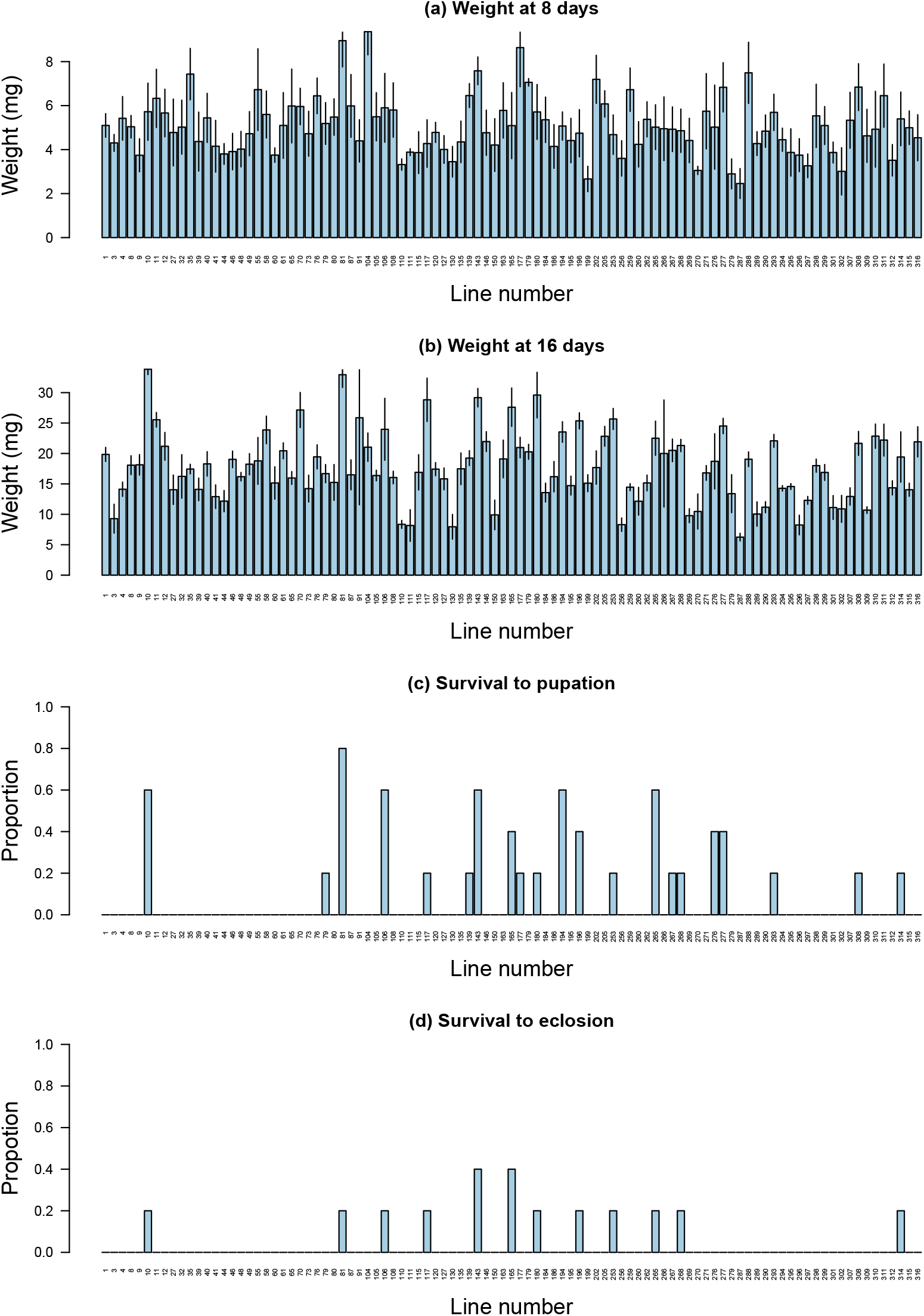
Barplots depict caterpillar performance traits as a function of *M. truncatula* inbred line. Colored bars denote the mean across replicates and vertical lines give the standard errors.

**Figure S4:**
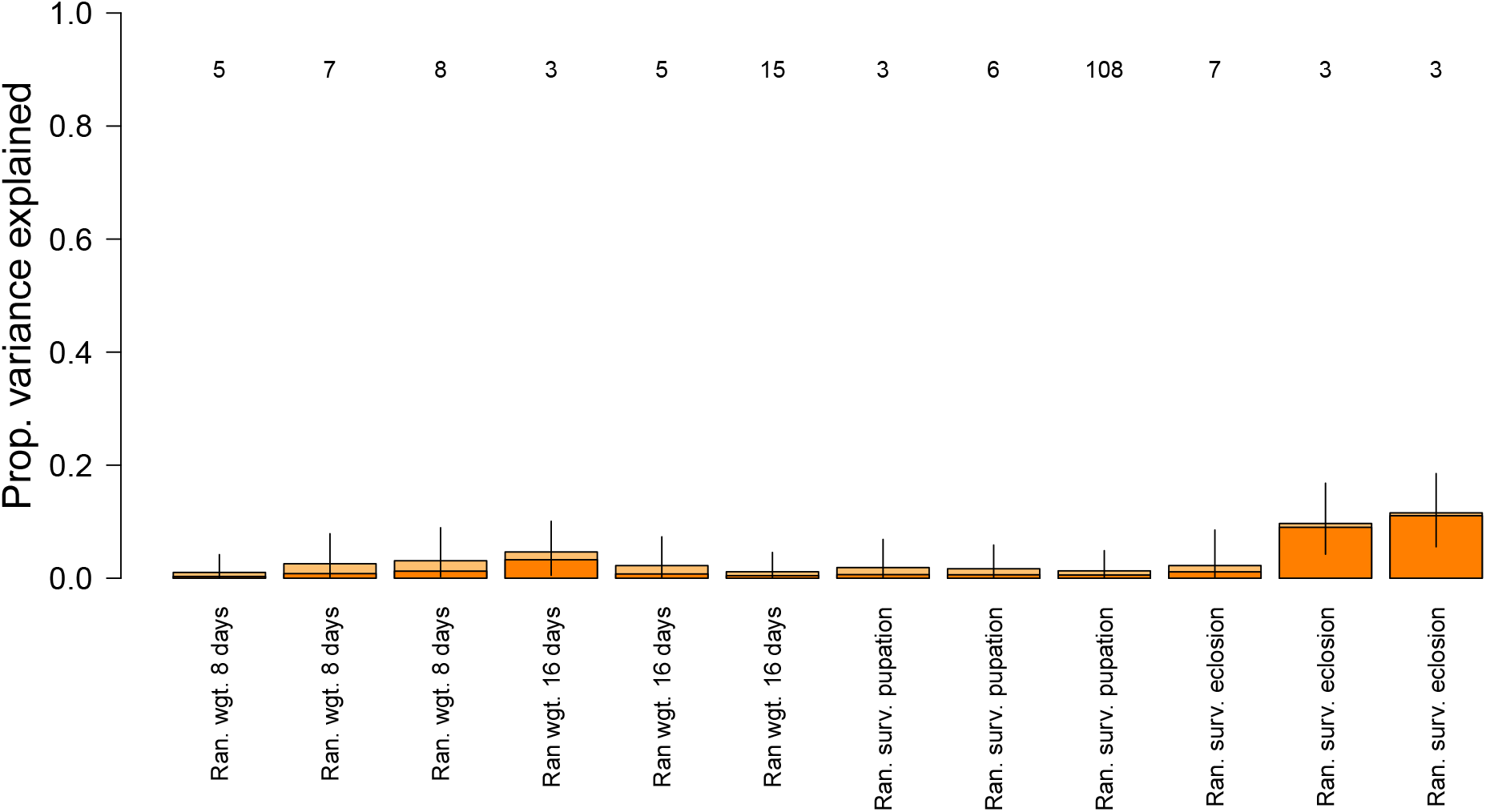
Graphical summary of permuted caterpillar (*L. melissa*) trait variation explained by *M. truncatula* genetics. Bars denote the posterior median for the proportion of permuted trait variation explained by plant genetics (PVE); vertical lines denote the 90% equal-tail probability intervals (ETPIs). Darker shaded regions of the bars provide a point estimate (posterior median) for the subset of the PVE that attributed to genetic variants with measurable effects (PGE; as opposed to infinitesimal effects). Numbers along the top of the plot give point estimates (posterior median) for the number of causal variants affecting each trait. Results are based on three replicate, randomized data sets where caterpillar trait data for each performance metric was permuted across *M. truncatula* lines (i.e., genotypes). Compare to results based on the true (unpermuted) data shown in Fig. 2.

**Figure S5:**
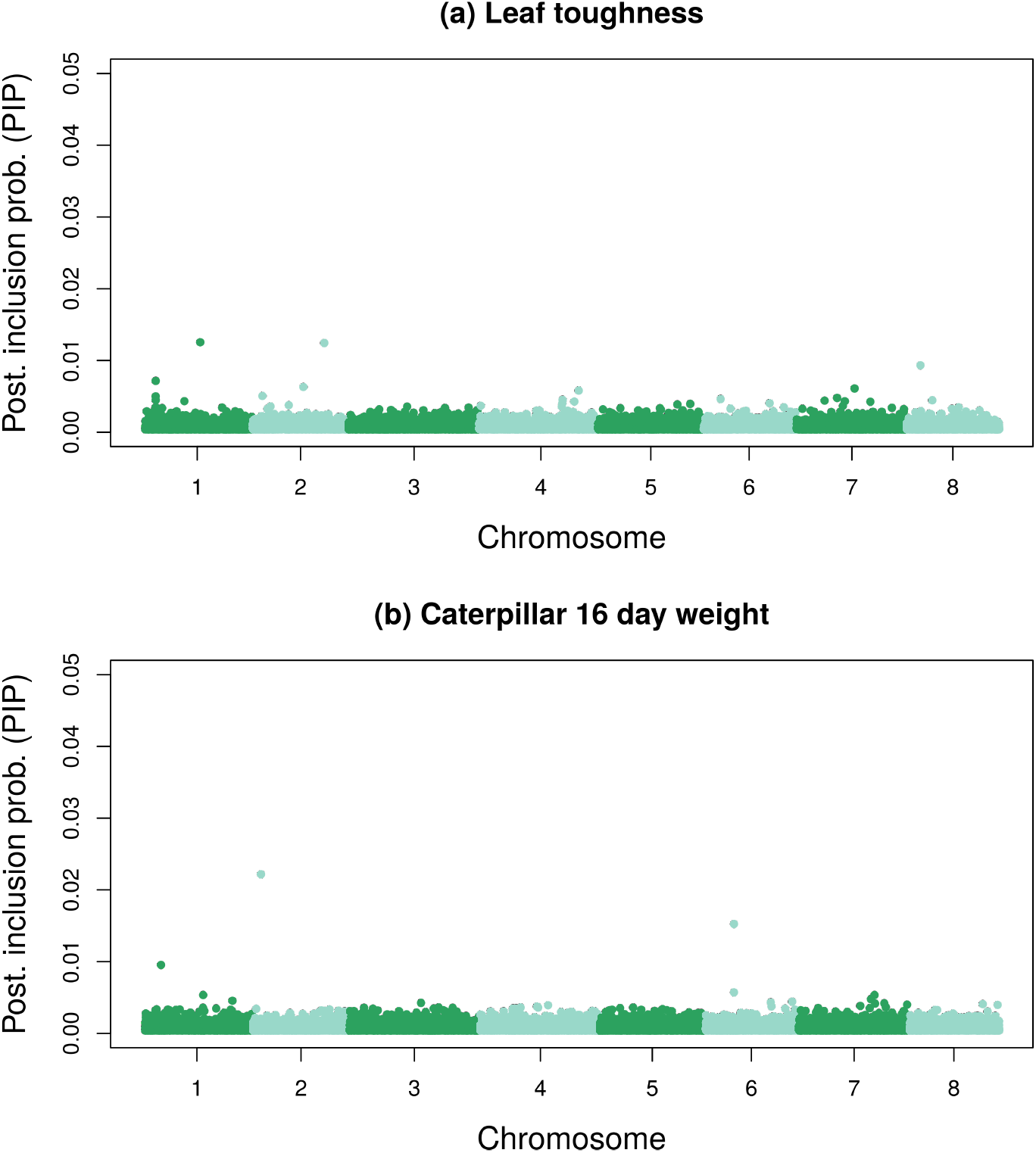
Manhattan plots showing posterior inclusions probabilities (PIPs) from Bayesian sparse linear mixed models relating (a) leaf toughness or (b) caterpillar weight at 16 days with *M. truncatula* genetics. The y-axis has been scaled to a PIP of 0.05 to show variability in PIPs among SNPs (no SNPs had PIPs higher than this for these traits). As with most traits we analyzed, individual SNPs do not have high PIPs; in other words, these are polygenic traits (compare to Fig. S6). Each point denotes a SNP, and SNPs are colored to show boundaries between chromosomes.

**Figure S6:**
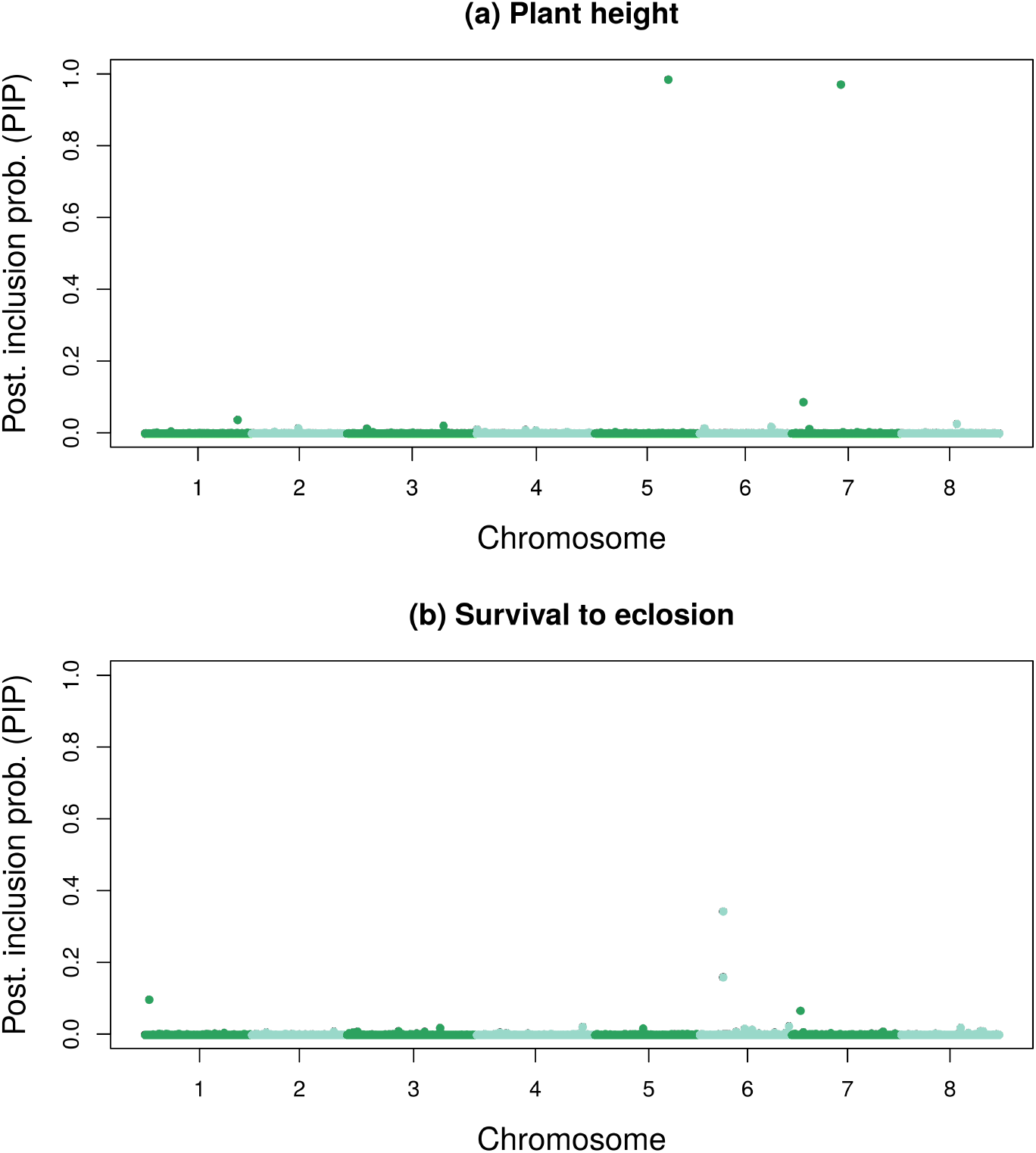
Manhattan plots showing posterior inclusions probabilities (PIPs) from Bayesian sparse linear mixed models relating (a) plant height or (b) survival to eclosion with *M. truncatula* genetics. These traits stand out as having individual SNPs with high PIPs (compare to Fig. S5). Each point denotes a SNP, and SNPs are colored to show boundaries between chromosomes.

**Figure S7:**
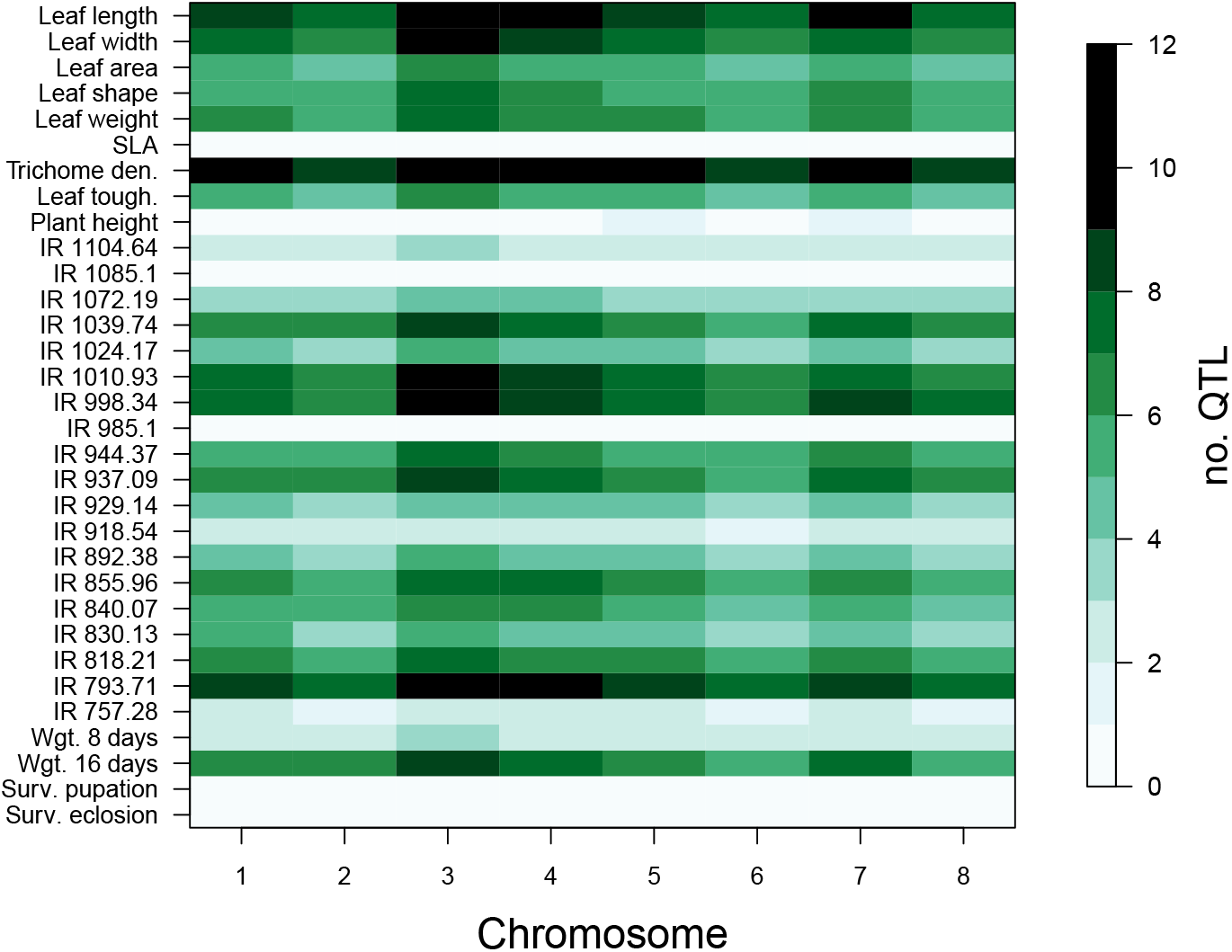
Heat map showing the number of QTL estimated for each trait on each of the eight *M. truncatula* chromosomes. The number of QTL per chromosome was estimated as the sum of the posterior inclusions probabilities across all SNPs on each chromosome. For most traits, the genetic signal (i.e., QTL) were spread uniformly across chromosomes (also see Fig. S5), but for a few traits, especially plant height and survival to eclosion, QTL were clustered on one or a few chromosomes (also see Fig. S6). Note that chromosome 3 is slightly larger than the other chromosomes and thus harbors a slight excess of QTL for most traits. The number of QTL per chromosome (and in general) also varies among traits. See Fig. 3 for the proportion of QTL for each trait on each chromosome.

**Figure S8:**
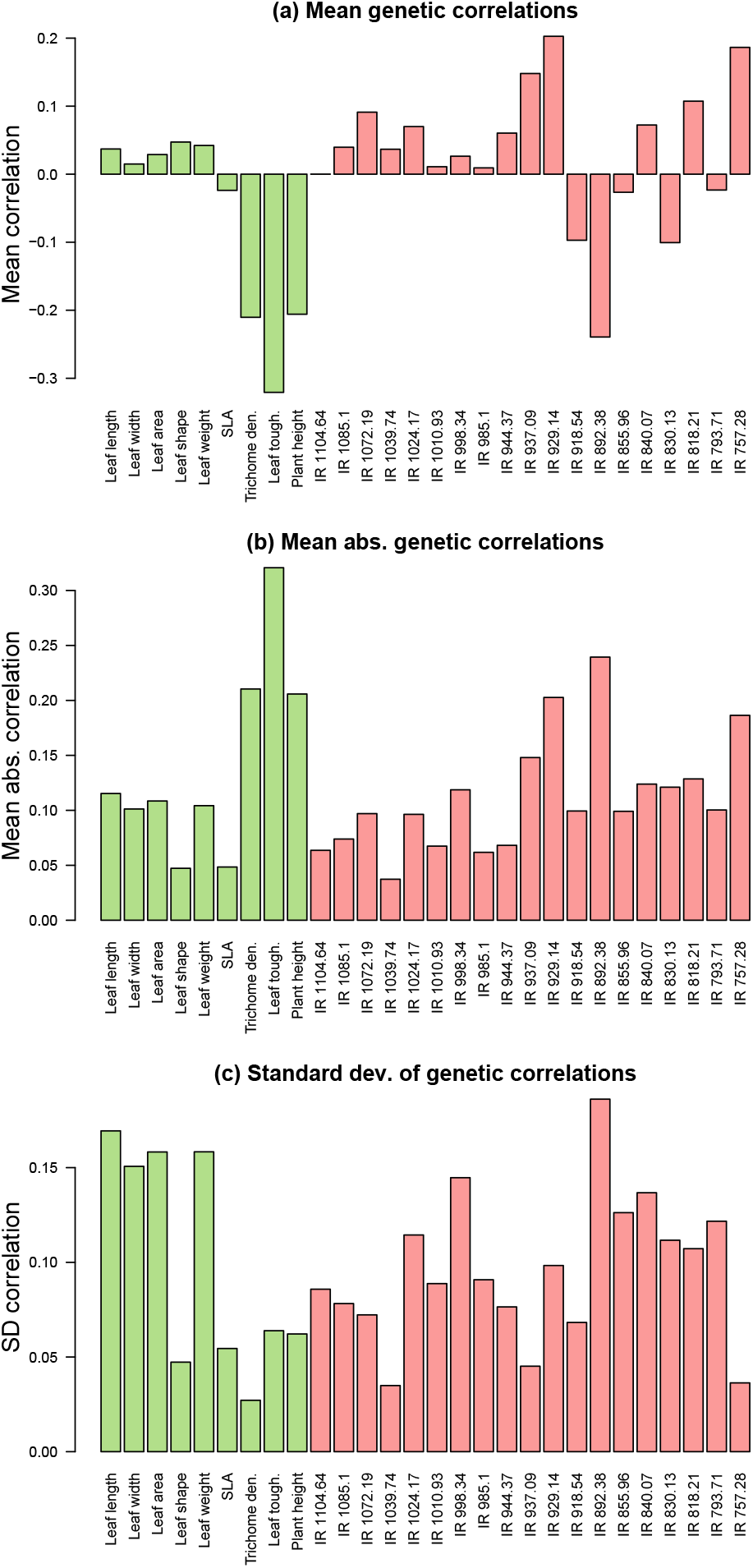
Barplots summarizing genetic correlations between plant traits and caterpillar performance traits. Panels (a) and (b) give the mean signed (a) or absolute value (b) genetic correlation between each plant trait and the four caterpillar performance traits. Panel (c) gives the standard deviation in genetic correlations across the four performance traits.

**Figure S9:**
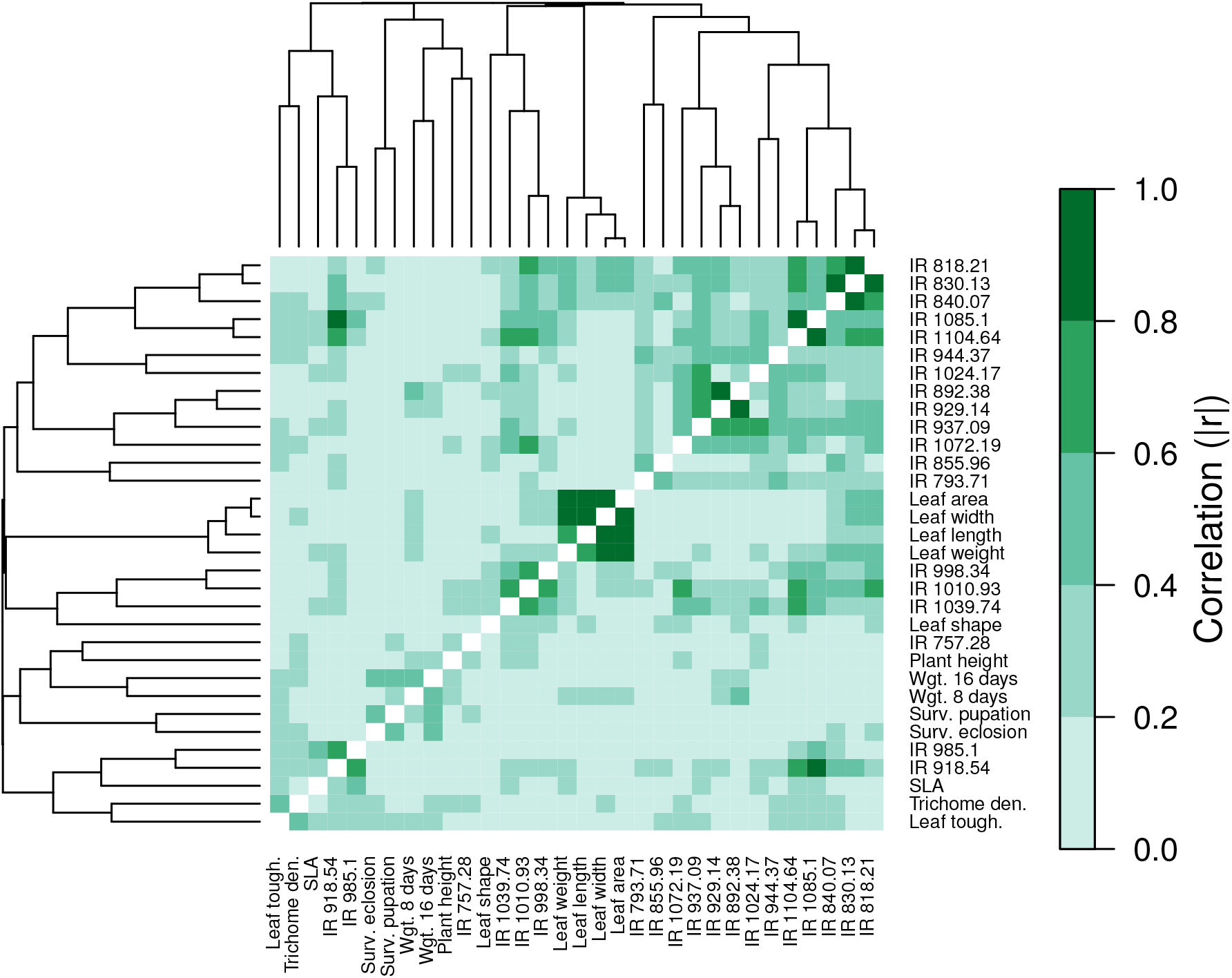
Heat map of genetic correlations for pairs of plant and caterpillar traits (this matrix is symmetric). Genetic correlations were computed from genomic estimated breeding values (GEBVs). Absolute values of correlations are shown. The dendrograms cluster traits by their genetic correlations and were computed with the `heatmap.2` function in `R`.

**Figure S10:**
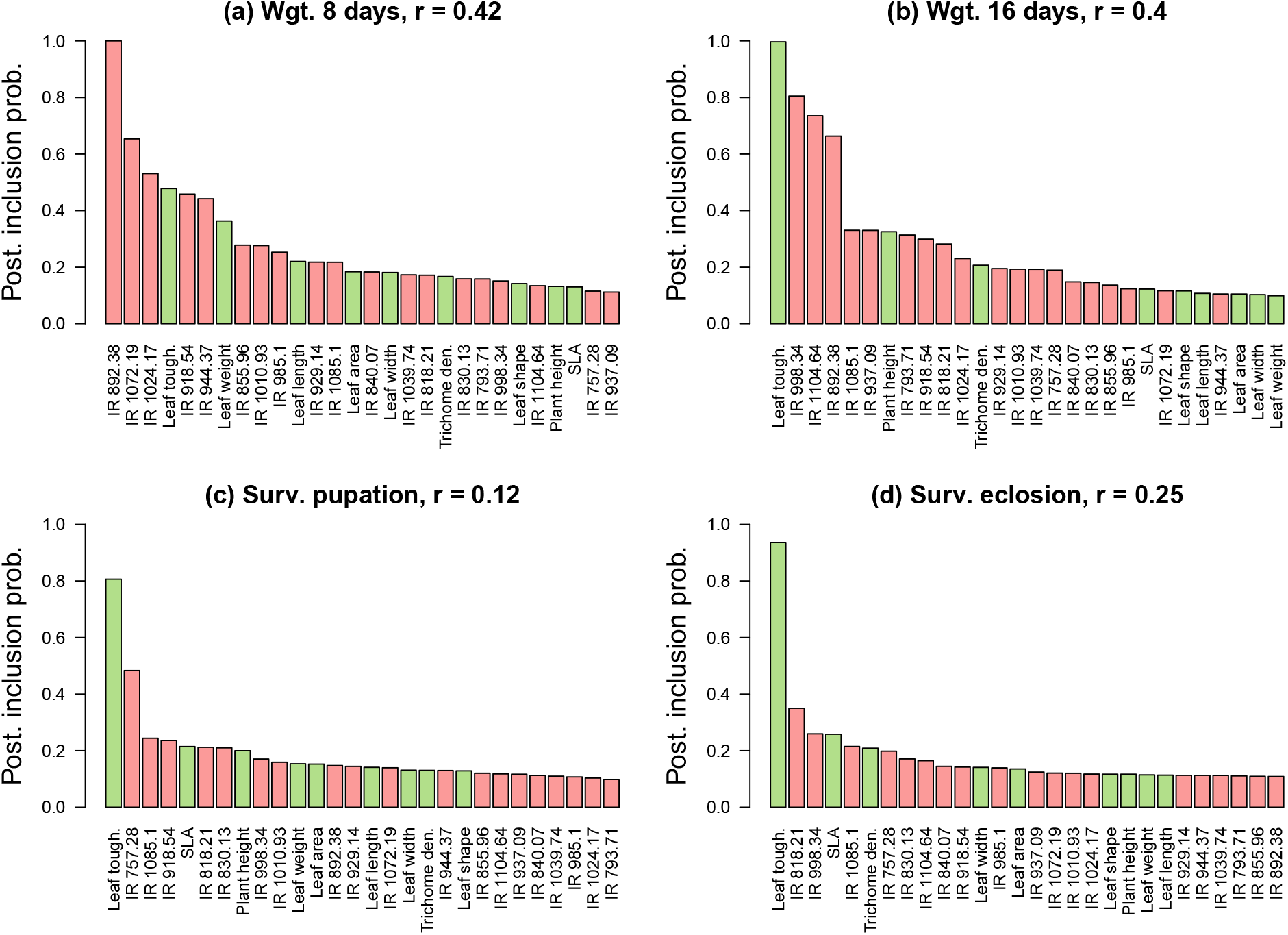
Barplots showing the effect of the genetic component of plant traits on the genetic component of caterpillar performance, specifically (a) weight at 8 days, (b) weight at 16 days, (c) survival to pupation, and (d) survival to eclosion. Bars denote Bayesian posterior inclusion probabilities (PIPs) for the effect of the genomic estimated breeding values (GEBVs) for each plant trait on the GEBVs for the caterpillar performance traits. Traits are sorted by their PIPs. Colors distinguish between plant growth/defense traits (green) and IR traits (pink). Pearson correlations between the caterpillar performance GEBVs and estimates of these from 10-fold cross-validation are given in the panel headers.

**Figure S11:**
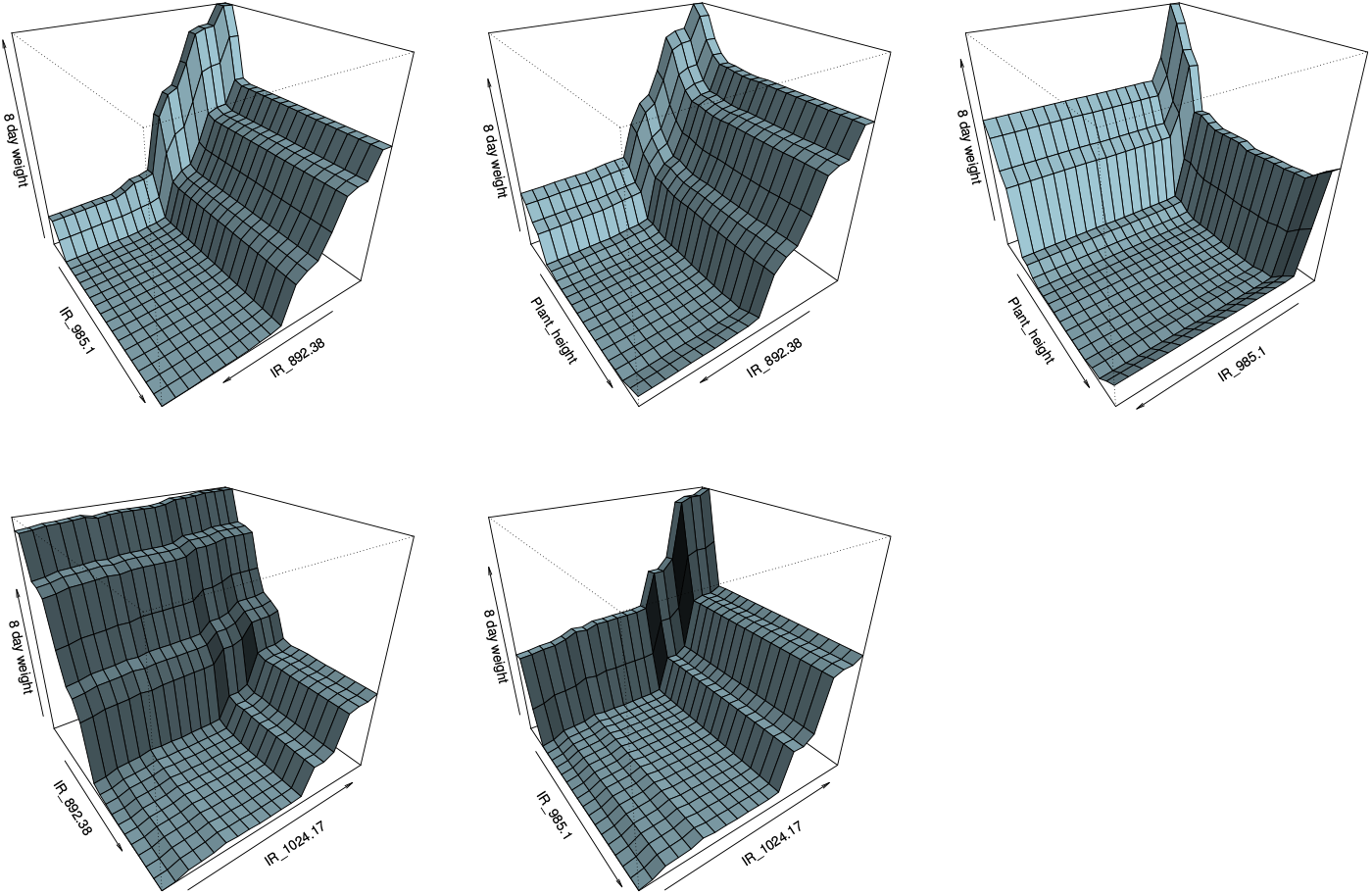
Graphical summary of interactions between pairs of plant trait (GEBVs) that best predict caterpillar 8-day weight GEBVs in the random forest regression analyses. Plots were generated in `plotmo` (Milborrow, 2018), and show interactions and relationships for the top traits.

**Figure S12:**
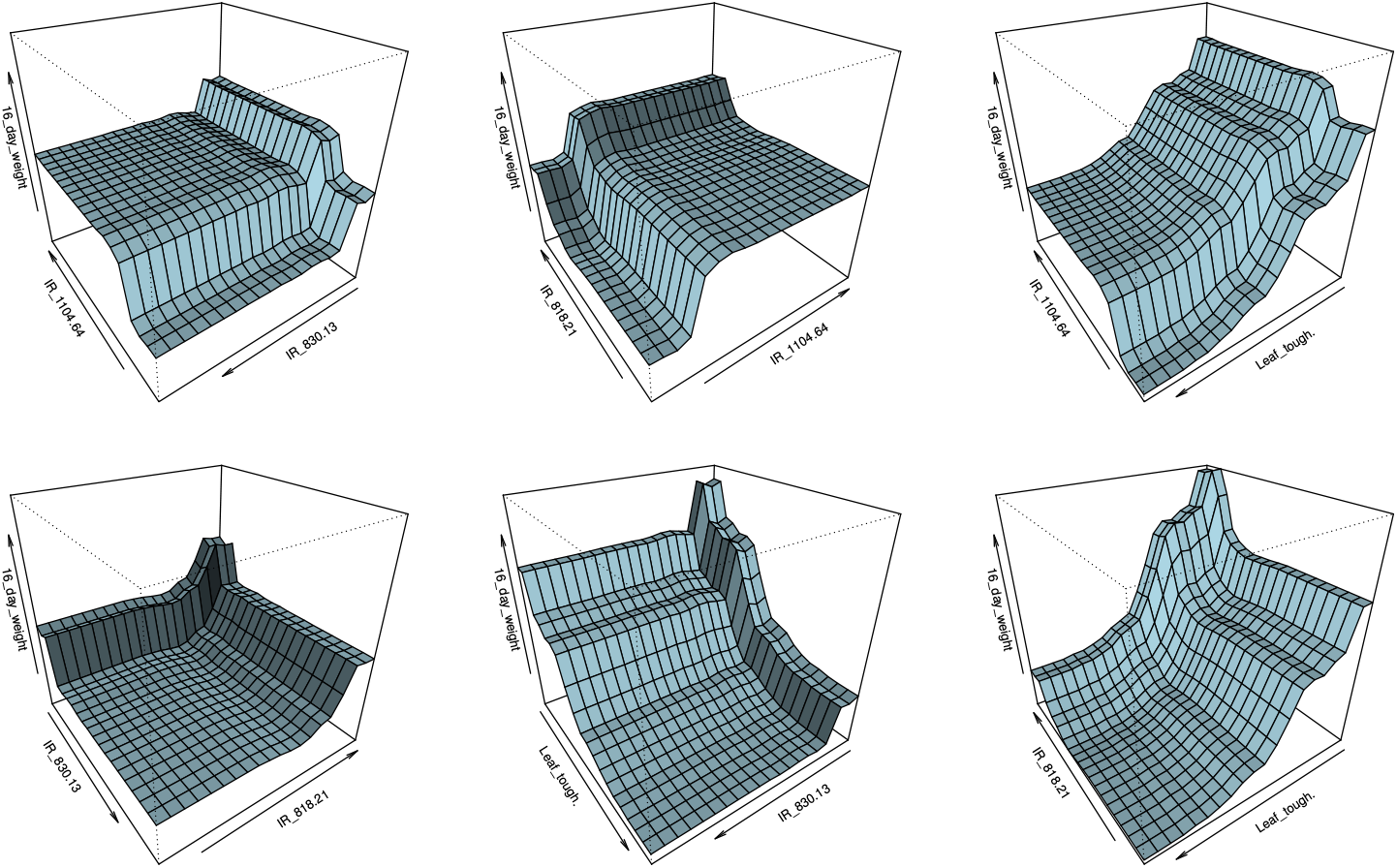
Graphical summary of interactions between pairs of plant trait (GEBVs) that best predict caterpillar 16-day weight GEBVs in the random forest regression analyses. Plots were generated in `plotmo` (Milborrow, 2018), and show interactions and relationships for the top traits.

## References

Allen HL, Estrada K, Lettre G, et al. (2010) Hundreds of variants clustered in genomic loci and biological pathways affect human height. Nature, 467, 832–838.

Bates D, Mächler M, Bolker B, Walker S (2015) Fitting linear mixed-effects models using lme4. Journal of Statistical Software, 67, 1–48.

Beavis WD (1998) QTl analyses: power, precision, and accuracy. In: Molecular Dissection of Complex Traits (ed. Patterson AH), pp. 145–162, CRC Press, New York.

Berenbaum MR, Zangerl AR (1998) Chemical phenotype matching between a plant and its insect herbivore. Proceedings of the National Academy of Sciences, 95, 13743–13748.

Bolnick DI, Svanbäck R, Fordyce JA, et al. (2002) The ecology of individuals: incidence and implications of individual specialization. The American Naturalist, 161, 1–28.

Braga MP, Guimaraes PR, Wheat CW, Soren N, Janz N (2018) Unifying host-associated diversification processes using butterflyplant networks. Nature Communications, 9, 5155.

Branca A, Paape TD, Zhou P, et al. (2011) Whole-genome nucleotide diversity, recombination, and linkage disequilibrium in the model legume *Medicago truncatula*. Proceedings of the National Academy of Sciences, 108, E864–E870.

Breiman L (2001) Random forests. Machine Learning, 45, 5–32.

Carius HJ, Little TJ, Ebert D (2001) Genetic variation in a host-parasite association: potential for coevolution and frequency-dependent selection. Evolution, 55, 1136–1145.

Carmona D, Lajeunesse MJ, Johnson MT (2011) Plant traits that predict resistance to herbivores. Functional Ecology, 25, 358–367.

Chaturvedi S, Lucas LK, Nice CC, Fordyce JA, Forister ML, Gompert Z (2018) The predictability of genomic changes underlying a recent host shift in Melissa blue butterflies. Molecular Ecology, 27, 2651–2666.

Choi HK, Kim D, Uhm T, et al. (2004a) A sequence-based genetic map of *Medicago truncatula* and comparison of marker colinearity with *M. sativa*. Genetics, 166, 1463–1502.

Choi HK, Mun JH, Kim DJ, et al. (2004b) Estimating genome conservation between crop and model legume species. Proceedings of the National Academy of Sciences of the United States of America, 101, 15289–15294.

Costa FR, Lang C, Almeida DR, Castilho CV, Poorter L (2018) Near-infrared spectrometry allows fast and extensive predictions of functional traits from dry leaves and branches. Ecological Applications.

Cox GW (2004) Alien species and evolution: the evolutionary ecology of exotic plants, animals, microbes, and interacting native species. Island Press.

Crainiceanu CM, Ruppert D (2004) Likelihood ratio tests in linear mixed models with one variance component. Journal of the Royal Statistical Society: Series B (Statistical Methodology), 66, 165–185.

Crutsinger GM, Collins MD, Fordyce JA, Gompert Z, Nice CC, Sanders NJ (2006) Plant genotypic diversity predicts community structure and governs an ecosystem process. Science, 313, 966–968.

Dambroski HR, Linn Jr C, Berlocher SH, Forbes AA, Roelofs W, Feder JL (2005) The genetic basis for fruit odor discrimination in *Rhagoletis* flies and its significance for sympatric host shifts. Evolution, 59, 1953–1964.

Danecek P, Auton A, Abecasis G, et al. (2011) The variant call format and vcftools. Bioinformatics, 27, 2156–2158.

Dawkins R (1982) The Extended Phenotype. Oxford University Press.

Edger PP, Heidel-Fischer HM, Bekaert M, et al. (2015) The butterfly plant arms-race escalated by gene and genome duplications. Proceedings of the National Academy of Sciences, p. 201503926.

Ehrlich PR, Raven PH (1964) Butterflies and plants: a study in coevolution. Evolution, 18, 586–608.

Eichler EE, Flint J, Gibson G, et al. (2010) Missing heritability and strategies for finding the underlying causes of complex disease. Nature Reviews Genetics, 11, 446–450.

Farkas TE, Mononen T, Comeault AA, Hanski I, Nosil P (2013) Evolution of camouflage drives rapid ecological change in an insect community. Current Biology, 23, 1835–1843.

Foley WJ, McIlwee A, Lawler I, Aragones L, Woolnough AP, Berding N (1998) Ecological applications of near infrared reflectance spectroscopy–a tool for rapid, cost–effective prediction of the composition of plant and animal tissues and aspects of animal performance. Oecologia, 116, 293–305.

Fordyce JA (2010) Host shifts and evolutionary radiations of butterflies. Proceedings of the Royal Society of London, Series B, 277, 3735–3743.

Forister ML, Dyer LA, Singer MS, Stireman III JO, Lill JT (2012) Revisiting the evolution of ecological specialization, with emphasis on insect–plant interactions. Ecology, 93, 981–991.

Forister ML, Gompert Z, Nice CC, Forsiter GW, Fordyce JA (2011) Ant association facilitates the evolution of diet breadth in a Lycaenid butterfly. Proceeding of the Royal Society B., 278, 1539–1547.

Forister ML, Nice CC, Fordyce JA, Gompert Z (2009) Host range evolution is not driven by the optimization of larval performance: the case of *Lycaeides melissa* (Lepidoptera: Lycaenidae) and the colonization of alfalfa. Oecologia, 160, 551–561.

Forister ML, Novotny V, Panorska AK, et al. (2015) The global distribution of diet breadth in insect herbivores. Proceedings of the National Academy of Sciences, 112, 442–447.

Forister ML, Scholl CF, Jahner JP, et al. (2013) Specificity, rank preference, and the colonization of a non-native host plant by the Melissa blue butterfly. Oecologia, 172, 177–188.

Gompert Z, Egan SP, Barrett RD, Feder JL, Nosil P (2017) Multilocus approaches for the measurement of selection on correlated genetic loci. Molecular Ecology, 26, 365–382.

Gompert Z, Fordyce JA, Forister ML, Nice CC (2008) Recent colonization and radiation of North American *Lycaeides* (*Plebejus*) inferred from mtDNA. Molecular Phylogenetics and Evolution, 48, 481–490.

Gompert Z, Jahner JP, Scholl CF, et al. (2015) The evolution of novel host use is unlikely to be constrained by trade-offs or a lack of genetic variation. Molecular Ecology, 24, 2777–2793.

Greven S, Crainiceanu CM, Küchenhoff H, Peters A (2008) Restricted likelihood ratio testing for zero variance components in linear mixed models. Journal of Computational and Graphical Statistics, 17, 870–891.

Guan Y, Stephens M (2011) Bayesian variable selection regression for genome-wide association studies and other large-scale problems. Annals of Applied Statistics, 5, 1780–1815.

Hanley ME, Lamont BB, Fairbanks MM, Rafferty CM (2007) Plant structural traits and their role in anti-herbivore defence. Perspectives in Plant Ecology, Evolution and Systematics, 8, 157–178.

Harrison JG, Gompert Z, Fordyce JA, et al. (2016) The many dimensions of diet breadth: Phytochemical, genetic, behavioral, and physiological perspectives on the interaction between a native herbivore and an exotic host. PloS ONE, 11, e0147971.

Harrison JG, Philbin CS, Gompert Z, et al. (2018) Deconstruction of a plant-arthropod community reveals influential plant traits with nonlinear effects on arthropod assemblages. Functional Ecology, 32, 1317–1328.

Hayes B, Bowman P, Chamberlain A, Goddard M (2009) Genomic selection in dairy cattle: Progress and challenges. Journal of Dairy Science, 92, 433–443.

Hendry AP (2016) Eco-evolutionary dynamics. Princeton University Press.

Kang Y, Sakiroglu M, Krom N, et al. (2015) Genome-wide association of drought-related and biomass traits with hapmap SNPs in *Medicago truncatula*. Plant, Cell & Environment, 38, 1997–2011.

Kennedy GG, Barbour JD (1992) Plant resistance to herbivores and pathogens: ecology, evolution, and genetics, chap. Resistance variation in natural and managed systems, pp. 13–41. University of Chicago Press, Chicago.

Lankau RA (2012) Coevolution between invasive and native plants driven by chemical competition and soil biota. Proceedings of the National Academy of Sciences, 109, 11240–11245.

Levin DA (1973) The role of trichomes in plant defense. The Quarterly Review of Biology, 48, 3–15.

Li H, Handsaker B, Wysoker A, et al. (2009) The Sequence Alignment/Map format and SAMtools. Bioinformatics, 25, 2078–2079.

Li W, Schuler MA, Berenbaum MR (2003) Diversification of furanocoumarin-metabolizing cytochrome P450 monooxygenases in two papilionids: specificity and substrate encounter rate. Proceedings of the National Academy of Sciences, 100, 14593–14598.

Liaw A, Wiener M (2002) Classification and regression by randomforest. R news, 2, 18–22.

Lucas LK, Nice CC, Gompert Z (2018) Genetic constraints on wing pattern variation in *Lycaeides* butterflies: A case study on mapping complex, multifaceted traits in structured populations. Molecular Ecology Resources, 18, 892–907.

Luijckx P, Ben-Ami F, Mouton L, Du Pasquier L, Ebert D (2011) Cloning of the unculturable parasite pasteuria ramosa and its *Daphnia* host reveals extreme genotype–genotype interactions. Ecology Letters, 14, 125–131.

Luijckx P, Fienberg H, Duneau D, Ebert D (2013) A matching-allele model explains host resistance to parasites. Current Biology, 23, 1085–1088.

Malishev M, Sanson GD (2015) Leaf mechanics and herbivory defence: how tough tissue along the leaf body deters growing insect herbivores. Austral Ecology, 40, 300–308.

Mandeville EG, Parchman TL, Thompson KG, et al. (2017) Inconsistent reproductive isolation revealed by interactions between catostomus fish species. Evolution Letters, 1, 255–268.

McKenna A, Hanna M, Banks E, et al. (2010) The Genome Analysis Toolkit: A MapReduce framework for analyzing next-generation DNA sequencing data. Genome Research, 20, 1297–1303.

Meuwissen THE, Hayes B, Goddard M (2001) Prediction of total genetic value using genome-wide dense marker maps. Genetics, 157, 1819–1829.

Michaud R, Lehman WF, Rumbaugh MD (1988) World distribution and historical developments. In: Alfalfa and Alfalfa improvement (eds. Hanson AA, Barnes DK, Hill RR), vol. 29, chap. World distribution and historical developments, Madison.

Milborrow S (2018) plotmo: Plot a Model’s Residuals, Response, and Partial Dependence Plots. R package version 3.5.1.

Mitchell C, Brennan RM, Graham J, Karley AJ (2016) Plant defense against herbivorous pests: exploiting resistance and tolerance traits for sustainable crop protection. Frontiers in plant science, 7, 1132.

Mitter C, Farrell B, Wiegmann B (1988) The phylogenetic study of adaptive zones: has phytophagy promoted insect diversification? The American Naturalist, 132, 107–128.

Moreau D (2006) Morphology, development and plant architecture of M. truncatula. In: The Medicago truncatula handbook, INRA.

Nallu S, Hill JA, Don K, et al. (2018) The molecular genetic basis of herbivory between butterflies and their host plants. Nature Ecology & Evolution, 2, 1418.

Nosil P (2004) Reproductive isolation caused by visual predation on migrants between divergent environments. Proceedings of the Royal Society of London Series B-Biological Sciences, 271, 1521–1528.

Nosil P, Villoutreix R, de Carvalho CF, et al. (2018) Natural selection and the predictability of evolution in *Timema* stick insects. Science, 359, 765–770.

Nouhaud P, Gautier M, Gouin A, et al. (2018) Identifying genomic hotspots of differentiation and candidate genes involved in the adaptive divergence of pea aphid host races. Molecular Ecology, 27, 3287–3300.

Nylin S, Agosta S, Bensch S, et al. (2018) Embracing colonizations: a new paradigm for species association dynamics. Trends in Ecology & Evolution, 33, 4–14.

Ober U, Ayroles JF, Stone EA, et al. (2012) Using whole-genome sequence data to predict quantitative trait phenotypes in *Drosophila melanogaster*. PLoS Genetics, 8, e1002685.

Ordas B, Malvar RA, Santiago R, Sandoya G, Romay MC, Butron A (2009) Mapping of QTL for resistance to the mediterranean corn borer attack using the intermated B73 × Mo17 (IBM) population of maize. Theoretical and Applied Genetics, 119, 1451–1459.

Purcell S, Neale B, Todd-Brown K, et al. (2007) PLINK: a tool set for whole-genome association and population-based linkage analyses. The American Journal of Human Genetics, 81, 559–575.

Ramirez JA, Posada JM, Handa IT, et al. (2015) Near-infrared spectroscopy (NIRS) predicts non-structural carbohydrate concentrations in different tissue types of a broad range of tree species. Methods in Ecology and Evolution, 6, 1018–1025.

Rausher MD, Simms EL (1989) The evolution of resistance to herbivory in *Ipomoea purpurea*. I. attempts to detect selection. Evolution, 43, 563–572.

Riesch R, Muschick M, Lindtke D, et al. (2017) Transitions between phases of genomic differentiation during stick-insect speciation. Nature Ecology & Evolution, 1, 0082.

Rockman MV (2012) The QTN program and the alleles that matter for evolution: All that’s gold does not glitter. Evolution, 66, 1–17.

Santure AW, Poissant J, De Cauwer I, et al. (2015) Replicated analysis of the genetic architecture of quantitative traits in two wild great tit populations. Molecular Ecology, 24, 6148–6162.

Scheipl F, Greven S, Kuechenhoff H (2008) Size and power of tests for a zero random effect variance or polynomial regression in additive and linear mixed models. Computational Statistics & Data Analysis, 52, 3283–3299.

Schoonhoven LM, van Loon JJA, Dicke M (2010) Insect-Plant Biology. 2nd edn., Oxford University Press.

Scott J (1986) The Butterflies of North America: A Natural History and Field Guide. Stanford University Press.

Soria-Carrasco V, Gompert Z, Comeault AA, et al. (2014) Stick insect genomes reveal natural selection’s role in parallel speciation. Science, 344, 738–742.

Stanton-Geddes J, Paape T, Epstein B, et al. (2013) Candidate genes and genetic architecture of symbiotic and agronomic traits revealed by whole-genome, sequence-based association genetics in *Medicago truncatula*. PloS ONE, 8, e65688.

Stowe KA (1998) Realized defense of artificially selected lines of *Brassica rapa*: effects of quantitative genetic variation in foliar glucosinolate concentration. Environmental Entomology, 27, 1166–1174.

Strauss SY, Lau JA, Carroll SP (2006) Evolutionary responses of natives to introduced species: what do introductions tell us about natural communities? Ecology Letters, 9, 357–374.

Thompson JN (2013) Relentless Evolution. University of Chicago Press.

Via S (1990) Ecological genetics and host adaptation in herbivorous insects: the experimental study of evolution in natural and agricultural systems. Annual Review of Entomology, 35, 421–446.

Vila R, Bell CD, Macniven R, et al. (2011) Phylogeny and palaeoecology of *Polyommatus* blue butterflies show Beringia was a climate-regulated gateway to the New World. Proceedings of the Royal Society B: Biological Sciences, 278, 2737–2744.

Weinig C, Stinchcombe J, Schmitt J (2003) QTL architecture of resistance and tolerance traits in *Arabidopsis thaliana* in natural environments. Molecular Ecology, 12, 1153–1163.

Weinig C, Ungerer M, Dorn L, et al. (2002) Novel loci control variation in reproductive timing in *Arabidopsis thaliana* in natural environments. Genetics, 162, 1875–1884.

Weiss KM (2008) Tilting at quixotic trait loci (QTL): an evolutionary perspective on genetic causation. Genetics, 179, 1741–1756.

Wen Z, Rupasinghe S, Niu G, Berenbaum MR, Schuler MA (2006) CYP6B1 and CYP6B3 of the black swallowtail (*Papilio polyxenes*): adaptive evolution through subfunctionalization. Molecular Biology and Evolution, 23, 2434–2443.

Wheat CW, Vogel H, Wittstock U, Braby MF, Underwood D, Mitchell-Olds T (2007) The genetic basis of a plant–insect coevolutionary key innovation. Proceedings of the National Academy of Sciences, 104, 20427–20431.

Whitham T, Bailey J, Schweitzer J, et al. (2006) A framework for community and ecosystem genetics: from genes to ecosystems. Nature Reviews Genetics, 7, 510–523.

Yang J, Benyamin B, McEvoy BP, et al. (2010) Common SNPs explain a large proportion of the heritability for human height. Nature Genetics, 42, 565–569.

Yoon S, Read Q (2016) Consequences of exotic host use: impacts on Lepidoptera and a test of the ecological trap hypothesis. Oecologia, 181, 985–996.

Young ND, Debellé F, Oldroyd GE, et al. (2011) The *Medicago* genome provides insight into the evolution of rhizobial symbioses. Nature, 480, 520.

Young ND, Udvardi M (2009) Translating *Medicago truncatula* genomics to crop legumes. Current Opinion in Plant Biology, 12, 193–201, genome Studies and Molecular Genetics.

Zellner A (1986) On assessing prior distributions and bayesian regression analysis with g-prior distributions. Bayesian inference and decision techniques.

Zeugner S, Feldkircher M (2015) Bayesian model averaging employing fixed and flexible priors: The BMS package for R. Journal of Statistical Software, 68, 1–37.

Zhou X, Carbonetto P, Stephens M (2013) Polygenic modeling with Bayesian sparse linear mixed models. PLoS Genetics, 9, e1003264.

